# Kappa Opioid–Oxytocin Interactions During Long-Term Partner Separation: Insights from PET Imaging in Titi Monkeys (Plecturocebus cupreus)

**DOI:** 10.64898/2026.07.30.739928

**Authors:** John P. Paulus, Claudia Manca, Alita J. D Almeida, Anelise Caceres, Meghan J. Sosnowski, Brad A. Hobson, Abhijit J. Chaudhari, Karen L. Bales

## Abstract

Social bonds are fundamental to health and well-being, and their disruption through partner separation is associated with significant physiological and psychological consequences. The kappa opioid receptor (KOR) system has been proposed as a key modulator of oxytocin (OT) release during partner separation, with prolonged separation hypothesized to trigger KOR downregulation in the paraventricular nucleus of the hypothalamus that disinhibits OT release over time (Bales & Rogers, 2022). However, this model has been difficult to assess non-invasively in primates. The present study used positron emission tomography (PET) with the KOR-selective radiotracer [^11^C]GR103545 to examine KOR availability *in vivo* across brain regions implicated in social bonding and separation distress in 16 pair-bonded titi monkeys using a within-subject two-week separation paradigm. Plasma OT, cerebrospinal fluid OT, and plasma cortisol were collected at each scan as complementary indices of peripheral and central OT signaling and physiological stress. We hypothesized that long-term separation would downregulate KORs in the hypothalamus and pituitary, indexed by reduced non-displaceable binding potential (BP_ND_), consistent with the Bales and Rogers model, and increase plasma OT consistent with KOR downregulation disinhibiting OT release. Partner separation significantly elevated plasma cortisol in both sexes, confirming the physiological stress of the manipulation. A significant Condition × Sex interaction was observed for plasma OT, reflecting a crossover pattern in which males, who had significantly higher plasma OT than females at baseline, showed a significant decrease during separation while females showed a non-significant increase, though the sex difference during separation did not reach significance. KOR availability was significantly reduced in the nucleus accumbens during separation, with a non-significant trend toward reduction in the anterior cingulate cortex, while no significant condition effects were observed in the remaining *a priori* chosen regions. No significant sex effects or Condition × Sex interactions were observed in any *a priori* chosen PET region. Together, these findings provide the first *in vivo* neuroimaging evidence of KOR system engagement during partner separation in a pair-bonded primate species, and reveal a sex-dependent OT response to separation that extends and adds nuance to the Bales and Rogers (2022) framework.

## Introduction

Social relationships are fundamental to human health and well-being. Strong social connections predict increased survival (Holt-Lunstad et al., 2010), and disruptions to these relationships carry health risks comparable to those of established risk factors such as cigarette smoking and obesity (Holt-Lunstad et al., 2010, 2015). In socially monogamous species, adults often form attachment relationships known as pair bonds that are characterized by a mutual preference for the partner (Bales et al., 2021). Healthy pair bonds are associated with improved immune and cardiovascular functioning, increased longevity, and positive effects on mood (Grewen et al., 2003; Kiecolt-Glaser & Wilson, 2017; Robles et al., 2014). The presence of a pair bonded partner can also act as a social buffer by reducing stress reactivity in humans (Ditzen et al., 2007; Heinrichs et al., 2003; Kirschbaum et al., 1995) and animals (Donovan et al., 2018; A. S. Smith & Wang, 2014; T. E. Smith et al., 1998). However, while proximity to a pair mate can reduce the negative effects of stressful stimuli, separation from a pair mate causes distress (Mason & Mendoza, 1998; McNeal et al., 2014; Mendoza & Mason, 1986; P. Sun et al., 2014). In humans, disruptions to the pair bond, such as divorce and bereavement, are associated with decreased neuroendocrine and immune functioning (Bower et al., 1998; Hopf et al., 2020; Irwin et al., 1988; Kiecolt-Glaser et al., 1987; Seiler et al., 2020).

The emotional and physiological consequences of partner separation are mediated by several interacting neurohormone systems. As a potent social stressor, partner separation activates the hypothalamic-pituitary-adrenal (HPA) axis, resulting in the release of corticotropin-releasing hormone (CRH) from the paraventricular nucleus (PVN) of the hypothalamus and downstream elevation of glucocorticoids such as cortisol and corticosterone. Specifically, elevated glucocorticoid concentrations, widely regarded as a physiological index of stress, have been observed across species during separation from a pair mate (Bosch et al., 2009; Hinde et al., 2016; Remage-Healey et al., 2003; P. Sun et al., 2014). Beyond its role in initiating the stress response, CRH has also been shown to directly modulate social behavior and neuropeptide release in regions central to pair bonding, including the nucleus accumbens (NAc) and PVN (Bosch et al., 2016; Pohl et al., 2019), suggesting that the HPA response to separation has consequences that extend well beyond peripheral stress hormone release.

Oxytocin (OT), a neuropeptide primarily synthesized in the PVN and supraoptic nuclei (SON) of the hypothalamus, plays a key role in reproduction and social behaviors (Borrow & Cameron, 2012). Intact social relationships are accompanied by increased brain OT signaling, which helps buffer against adverse life experiences (Carter et al., 2020; A. S. Smith & Wang, 2014), in part through the ability of a bonded partner’s presence to stimulate OT release and attenuate HPA axis reactivity. In the context of pair bonds, OT interacts with the reward system to facilitate both the formation and maintenance of the bond (Liu & Wang, 2003; Loth & Donaldson, 2020). The NAc, a key region for social reward, has garnered particular interest given its established role in mediating the rewarding effects of social contact and its sensitivity to OT signaling during pair bond formation and maintenance (Bosch et al., 2016; Bosch & Young, 2018; Insel & Shapiro, 1992).

Given its role in social bonds, OT levels are affected in partner loss. However, the direction and magnitude of OT changes appear to depend critically on the duration of separation and the sex of the individual. In prairie voles, short-term separation (3 days) from a partner decreases OT mRNA in the PVN and OTR binding in the NAc (Bosch et al., 2016), suggesting an initial suppression of the OT system. In contrast, following long-term separation (4 weeks), there is an increase in OT-immunoreactive cells in the PVN (P. Sun et al., 2014), suggesting a compensatory upregulation of OT over time. In humans, elevated plasma OT has been observed in the context of relationship distress, with evidence suggesting this response may be more pronounced in women than in men (Taylor et al., 2010).

As a socially monogamous primate species, titi monkeys offer a valuable translational model for investigating the neurobiology of separation, bridging rodent and human studies. Prior work in titi monkeys has examined the role of mu and kappa opioid receptors in pair bonds and social behavior, providing important groundwork for the present study (Ragen et al., 2013, 2015). In both short-term (2 days) and long-term (2 weeks) separation, cerebrospinal fluid (CSF) OT levels were elevated in male titi monkeys, while plasma OT was unaffected by separation and only increased upon reunion with the partner (Hinde et al., 2016). These findings highlight the importance of distinguishing between central and peripheral OT responses to separation and underscore the need for complementary *in vivo* neuroimaging approaches that can capture the broader neuromodulatory context in which OT changes occur.

While CRH and OT are the two neurohormone systems most consistently implicated in the response to partner separation, they do not act independently. OT has been shown to facilitate the return of the HPA axis to homeostatic levels following stress (Love, 2018), while conversely, activation of the Corticotropin-releasing hormone receptor 2 (CRHR2) receptors inhibits OT neurons in the PVN and reduces OT release in the NAc (Bosch et al., 2016; Pohl et al., 2019), suggesting a bidirectional relationship between the stress and social bonding systems. However, changes in CRH are not always correlated with changes in OT during separation (Bosch et al., 2009), indicating that the relationship between these two systems is not straightforward and that additional modulatory mechanisms may be involved. The observation that separation produces duration-dependent and sex-dependent changes in OT that are not fully explained by CRH activity alone points to the existence of an intermediary system capable of modulating OT release in a context-sensitive manner. The kappa opioid system is a compelling candidate for this role, given its established interactions with both the stress response and the oxytocinergic system, and its well-documented involvement in the negative affective states associated with social loss.

Opioids have long been implicated in social behavior (Panksepp et al., 1980). While mu opioid receptor (MOR) activation supports the rewarding, affiliative aspects of social contact and is necessary for initial pair bond formation (Resendez et al., 2013), kappa opioid receptor (KOR) activation produces the opposite effect, generating feelings of dysphoria and aversion across a range of contexts including stress (Callaghan et al., 2018) and drug withdrawal (Bruchas et al., 2010). The primary endogenous ligand for KORs is dynorphin, which is released in response to stress (Bruchas et al., 2010; Land et al., 2008). Importantly, the KOR system has been specifically implicated in behavioral responses to social stress, with endogenous dynorphin/KOR activation mediating the aversive behavioral consequences of social defeat in rodents (McLaughlin et al., 2006; Williams et al., 2018), and KOR signaling in the dorsal raphe producing sex-dependent effects on social behavior following social stress (Wright et al., 2018). In the context of pair bonding, KOR activity, particularly in the NAc, promotes behaviors necessary for pair bond maintenance, including mate-guarding and selective aggression toward novel conspecifics (Resendez et al., 2012). Critically, dynorphin/KOR signaling has been shown to inhibit OT release from the posterior pituitary, with KOR agonists decreasing and KOR antagonists increasing circulating OT concentrations in rodents (Summy-Long et al., 1990; Van de Heijning et al., 1991; van Wimersma Greidanus et al., 1996). This inhibitory effect is further connected to the stress response through the finding that dynorphin release is triggered by CRH receptor activation during stress (Land et al., 2008), providing a mechanistic pathway through which the HPA axis response to stress can suppress peripheral OT release via the KOR system.

Together, these observations position the KOR system as a plausible neuromodulator of OT release during partner separation, capable of translating the stress of social loss into downstream changes in oxytocinergic signaling. Despite this compelling theoretical framework, no study has directly examined KOR availability *in vivo* during partner separation in any species. While recent work has established the feasibility of [^11^C]GR103545 in positron emission tomography (PET) imaging for characterizing KOR binding in titi monkeys (Almeida et al., 2025), and has recently examined KOR availability in the context of acute stress and social buffering in this species (Manca et al., 2026), the application of this approach to directly test the role of KORs during long-term partner separation represents a critical gap in the literature that the present study addresses.

Sex differences in both the KOR and OT systems are well documented and are likely to shape the neurobiological response to partner separation in a sex-dependent manner. With respect to KOR, males and females differ in their sensitivity to KOR activation, with sex differences in KOR binding observed across multiple brain regions (Vijay et al., 2016). In the context of social behavior specifically, KOR activity in the NAc plays a sex-specific role in pair bond maintenance in prairie voles (Resendez et al., 2016). With respect to OT, females generally show greater OT reactivity to social stressors than males, consistent with the stress response described in humans (Taylor et al., 2000), and sex differences in OTR distribution across brain regions further contribute to divergent OT responses between the sexes (Dumais & Veenema, 2016). Together, these findings suggest that males and females may engage the KOR and OT systems differently in response to partner separation, potentially producing divergent neuroendocrine and neurochemical profiles during prolonged social loss.

Based on the converging evidence outlined above, Bales and Rogers (2022) proposed an expanded neurobiological model of partner loss that incorporates the KOR system as a key modulator of OT release during separation. Consistent with the original model proposed by Pohl et al. (2019), partner separation activates the HPA axis and triggers CRH release from the PVN. The expanded model adds that CRH-induced activation of CRHR2 receptors stimulates dynorphin release, leading to increased KOR activity in oxytocinergic neurons of the PVN in the NAc. During short-term separation, this KOR activation inhibits OT release in the NAc, contributing to the dysphoric, aversive state associated with acute partner loss. However, with prolonged separation, the model proposes that KORs in the PVN undergo downregulation, relieving the inhibitory brake on OT neurons and allowing central and peripheral OT levels to increase over time. This elevation of OT during long-term separation has theoretical plausibility from a functional standpoint, given OT’s established role in promoting social motivation and facilitating new bond formation (Gordon et al., 2011). Elevated OT during prolonged separation may serve to prime the social affiliation system for renewed bonding, whether with the original partner upon reunion or with a new conspecific if the original partner remains unavailable. Concurrently, KOR activity in the NAc shell is predicted to remain elevated or increase, sustaining the aversive motivational state that drives reunion-seeking behavior. The specific predictions of this model, particularly the proposed sex differences in OT response and the regional KOR dynamics in the hypothalamus and NAc, have yet to be directly evaluated using *in vivo* neuroimaging in any pair-bonded species.

The present study used PET imaging with the KOR-selective radiotracer [^11^C]GR103545 to examine *in vivo* KOR availability across brain regions implicated in social bonding, separation distress, and KOR-OT interactions in a sample of socially monogamous subjects scanned under two conditions: a baseline condition in which subjects had been living with their pair mate for at least two weeks, and a separation condition following two weeks of partner separation. This within-subject design allowed each subject to serve as their own control, maximizing statistical sensitivity to condition-related changes in KOR availability while controlling for individual differences in baseline receptor binding. PET with [^11^C]GR103545 allows for *in vivo* quantification of KOR availability, indexed as non-displaceable binding potential (BP_ND_) across brain regions with the cerebellum as the reference region, with decreases in BP_ND_ reflecting either increased endogenous dynorphin occupancy or receptor downregulation, both of which are relevant to the predictions of the expanded model of separation (Bales & Rogers, 2022). To complement the PET data, plasma OT, CSF OT, and plasma cortisol were measured at each scan as indices of peripheral and central OT signaling and physiological stress, respectively. Sex was included as a between-subjects variable in all analyses given the well-documented sex differences in both the KOR and OT systems described above. Based on the Bales and Rogers (2022) model and the broader literature reviewed here, we hypothesized that: (1) partner separation would elevate cortisol, confirming the physiological stress of the manipulation; (2) long-term separation would be associated with reduced KOR availability (BP_ND_) in the hypothalamus and pituitary, consistent with the Bales and Rogers (2022) model predicting KOR downregulation in the PVN as the system adapts to prolonged social loss, and with changes in NAc BP_ND_ reflecting KOR system engagement during the aversive motivational state of separation; (3) plasma OT would increase during separation relative to baseline, consistent with the long-term separation phase of the expanded model, and this effect would be more pronounced in females than males given their greater OT reactivity to social stressors.

## METHODS

### Subjects

Data were collected from 16 pair-bonded (8 males, 8 females) adult coppery titi monkeys (*Plecturocebus cupreus*) housed at the National Biomedical Research Institute (NBRI). Subjects were non-reproductive but gonadally intact, as the males had undergone vasectomy. The mean age of the subjects at the start of the study was 8.3 years old (range: 5-16y). All subjects had been paired with their mate at least two years prior to the study, with the mean duration of pairing being 4.8 years (range: 2-8y). The animals were housed in cages measuring 1.2 m x 1.2 m x 2.1 m or 1.2 m x 1.2 m x 1.8 m, under conditions similar to those described in Mendoza & Mason (1986) and Tardif et al. (2006). Rooms were maintained at 21°C on a 12:12-h light:dark cycle with lights on at 0600 and lights off at 1800. All animals were fed daily at 0800 and 1300 h on a diet consisting of monkey chow, carrots, apples, bananas, and rice cereal. Water was available ad libitum. During the separation period, subjects were housed in rooms maintained under the same environmental conditions as their home rooms. All experimental procedures were in agreement with the Guide for the Care and Use of Laboratory Animals and approved by the Institutional Animal Care and Use Committee (IACUC; protocol number #23483) of the University of California, Davis.

### Experimental Design

A within-subjects design was employed to examine the neurobiological effects of long-term partner separation in adult titi monkeys. Each subject underwent two social conditions: (1) a baseline (paired) condition and (2) a long-term separation condition. At the end of each social condition, subjects were anesthetized and underwent a [^11^C]GR103545 PET scan to assess changes in KOR availability across brain regions implicated in social bonding and separation distress (see Figure 1). The order of conditions was counterbalanced across subjects to minimize potential order effects, with a minimum interval of 28 days (average: 98 days) between successive scans to allow sufficient time for neural and hormonal measures to return to baseline. Because the experimental procedures involved separating a focal subject from their pair mate, each subject experienced two separation events over the course of the study: one as the focal subject (in which they remained in the home cage while the pair mate was relocated) and one as the partner (in which they were the one relocated while their pair mate served as the focal subject). The 28-day minimum interval applied between any reunification of the pair and the next scan, regardless of which member of the pair was being scanned, to ensure that both members of the pair had sufficient time to recover from their respective separation experiences before the next condition began.

**Figure 1.**
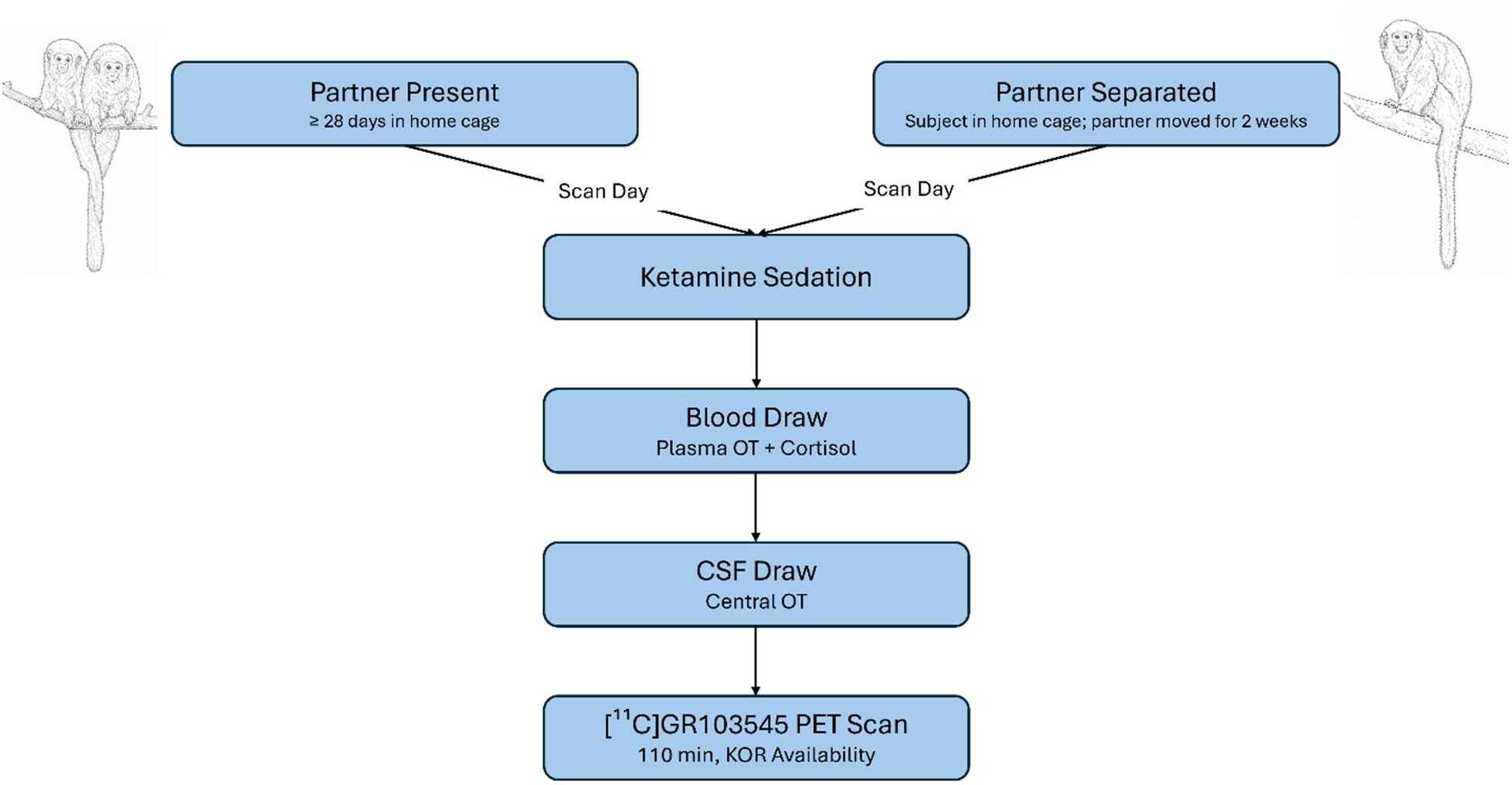
Experimental design. Subjects underwent two within-subject conditions in counterbalanced order with a minimum interval of 28 days between scans (mean interval: 98 days). In the baseline condition, subjects remained with their pair mate in the home cage for at least 28 days prior to the scan. In the separation condition, subjects remained in the home cage while their pair mate was relocated to a separate room for two weeks to eliminate visual and auditory contact. On the scan day for both conditions, subjects were sedated with ketamine, after which a blood sample was collected for measurement of plasma oxytocin and cortisol, followed by a CSF sample for measurement of central oxytocin. Subjects then underwent a 110-minute [^11^C]GR103545 PET scan to assess KOR availability across brain regions implicated in social bonding and separation distress. OT = oxytocin; CSF = cerebrospinal fluid; KOR = kappa opioid receptor.

During the baseline condition, subjects remained undisturbed with their established pair mate in their home cage for at least four weeks prior to the scan to ensure stable social bonds. For the long-term separation condition, subjects were separated from their pair mate for two weeks, a duration previously shown to elicit measurable behavioral and neurochemical effects in titi monkeys (Hinde et al., 2016). To avoid confounding the social stress of separation with the stress of a novel environment, the subject remained in the home cage while the pair mate was moved to a different room to eliminate visual and auditory access to their partner. Subjects retained visual and auditory access to other titi monkeys housed in the same room throughout the separation period, ensuring that any observed effects reflected the specific loss of the pair mate rather than complete social isolation.

Prior to each PET scan, subjects were removed from their home cage and sedated. A blood sample (1 ml) was collected from the femoral vein using heparinized syringes for measurement of plasma OT and cortisol, and a CSF sample (∼200 µl) was collected via cisternal puncture for measurement of central OT. The mean time from initial handling to blood collection was 6 minutes 40 seconds across both conditions (baseline: 7 minutes 1 second; separation: 6 minutes 38 seconds). Following sample collection, subjects were prepared for PET imaging as described below.

### PET Imaging Procedures

PET imaging was performed using the KOR-selective radioligand [¹¹C]GR103545, which has been characterized in humans (Naganawa et al., 2014) and more recently in titi monkeys (Almeida et al., 2025). Two intravenous catheters were placed in the saphenous veins of both legs to administer IV fluids (lactated Ringers solution, 10 ml/kg/h), and an endotracheal tube was inserted to maintain a clear airway and allow for monitoring of ventilation and aspiration.

Subjects were positioned head-first supine within a dedicated brain PET scanner (PiPET, Brain Biosciences, Rockville, MD) and imaged for 110 minutes. The [^11^C]GR103545 radiotracer was synthesized by the UC Davis Center for Molecular and Genomic Imaging using established methods (Nabulsi et al., 2011). PET acquisition began approximately 15 seconds prior to iv injection of [^11^C]GR103545 (injected activity 47.7 ± 5.7 MBq). A single-pass cold transmission scan was acquired pre-injection for attenuation and scatter correction. Data were reconstructed with an isotropic voxel size of 0.8 mm^3^ using framing of 6 x 10 s, 8 x 30 s, 5 x 60 s, 4 x 300 s, 8 x 600 s. Anesthesia was maintained throughout the scan with isoflurane (Cat. No. 66794-017, 1-2%, Medline Medical). The mean time from sedation to PET scan start was 64 minutes across both conditions (baseline: 64 minutes 43 seconds; separation: 64 minutes).

Following the scan, subjects were temporarily housed individually in a metabolism room for post-scan survey, after which they were returned to their home cage and reunited with their pair mate.

### Hormone Analyses

Upon collection, blood and CSF samples were immediately placed on ice. Blood samples were centrifuged at 3000 RPM for 15 minutes at 4°C, after which the plasma was separated and transferred into two sets of tubes for OT and cortisol assay separately. Plasma and CSF samples were stored at -80°C until assay. CSF samples were assayed for OT and plasma samples were assayed for both OT and cortisol. Plasma OT and CSF OT were measured using a commercially available enzyme immunoassay kit (Cat. No. K048-H; ELISA; Arbor Assays, Ann Arbor, MI, USA) that was validated for use with titi monkey CSF and plasma samples. Samples were run in duplicate when sufficient volume was collected.

CSF samples were not extracted prior to assay, as per previous literature suggesting that CSF samples do not require extraction (Dal Monte et al., 2014; Parker et al., 2010; Tabak et al., 2023). Out of 32 CSF collections, 19 samples did not meet the minimum required volume or fell below the assay detection threshold, resulting in 13 valid CSF observations available for analysis. Of the 11 samples run in duplicate, the intra-assay coefficient of variation (CV) was 5.05%.

Plasma OT samples were extracted prior to assay using a reduction/alkylation procedure (see Supplementary Materials), consistent with previous literature indicating that extraction is necessary for plasma to yield valid results (Szeto et al., 2011; Tabak et al., 2023). Plasma OT samples were all run in duplicate across two plates. The intra-assay CVs were 5.78% for plate 1 and 2.12% for plate 2. Because pooled quality control samples were not run across both plasma OT plates, inter-assay reliability was assessed using Standard 4, which falls within the dynamic range of the standard curve where the assay is most reliable. The inter-assay CV for plasma OT was 1.72%. Out of 32 blood sample collections, 3 did not meet the minimum required volume or fell below the assay detection threshold, resulting in 29 valid plasma observations available for analysis.

Plasma cortisol was assayed at the UC Davis Endocrinology Laboratory using commercial radioimmunoassay kits (Siemens Healthcare, Malvern, PA, USA), previously validated both chemically and biologically for use with titi monkey plasma samples (Witczak et al., 2021). Thermo Nunc Maxisorp plates were coated overnight at 4°C with cortisol antisera (R4866; UC Davis Endocrinology lab). Plasma samples were not extracted but were diluted 1:1000 in EIA buffer (0.1 M phosphate buffer, pH 7.0, 0.1% BSA). Samples and standards (0.98-500 pg/well) were incubated with cortisol-HRP conjugate overnight at 4°C. Following washing, plates were developed using ABTS substrate and read at 405/570 nm. Samples were run in duplicate across two plates, with intra-assay CVs of 13.5% and 2.9% for plates 1 and 2 respectively, and an inter-assay CV of 1.2%.

### MRI Acquisition

Structural MRI scans were acquired to provide anatomical reference images for PET data co-registration and region of interest delineation. Scans were conducted on a separate day from PET imaging using a GE Signa LX 9.1 scanner (General Electric Corporation, Milwaukee, WI, USA) with a 1.5 T field strength and a 3-inch surface coil. Subjects were fasted for 8 to 12 hours prior to the procedure and sedated with ketamine (10 mg/kg IM) and midazolam (0.1 mg/kg IM), and an endotracheal tube was placed. Physiological parameters including EtCO2, oxygen saturation, blood pressure, and heart rate were monitored throughout the scan. T_1_-weighted anatomical images were acquired using a 3D spoiled gradient echo pulse sequence in the coronal plane with the following parameters: echo time (TE) = 7.9 ms, repetition time (TR) = 22.0 ms, flip angle = 30.0°, field of view = 8 cm, number of excitations = 3, matrix size = 256 × 256, resulting in a pixel size of 0.3125 × 0.3125 mm^2^ and a slice thickness of 1 mm. For subjects with an existing MRI acquired within the same year as the start of the study, the previously acquired scan was used in place of a new acquisition.

### Image Analysis

PET and MRI data were processed using a standardized image analysis pipeline developed specifically for titi monkey neuroimaging data (Almeida et al., 2026). Each subject’s MR image data were resampled to an isotropic voxel size of 0.3 mm^3^, bias corrected, and skull stripped. Scans were then registered to a titi monkey brain reference space using the symmetric normalization algorithm implemented in the Advanced Normalization Tools (ANTs) toolkit (Avants et al., 2011), and a titi monkey brain atlas was warped to each individual MRI to generate subject-specific regional labels (Almeida et al., 2026). Reconstructed PET images were imported into PMOD software (version 4.5, PMOD Technologies, Zurich, Switzerland), where each scan was visually inspected for data quality, motion corrected, and manually co-registered to the corresponding anatomical MRI. Atlas-derived regional labels were then transferred to the co-registered PET data to extract region-specific time-activity curves. Non-displaceable binding potential (BP_ND_) was calculated for each region of interest using the Simplified Reference Tissue Model (SRTM), with the cerebellum serving as the reference region, consistent with prior applications of this radiotracer in titi monkeys and other non-human primates (Almeida et al., 2025). Figure 2 shows representative MRI, PET, and overlay images from a single subject, illustrating the typical data quality and anatomical coverage achieved with this imaging pipeline.

**Figure 2.**
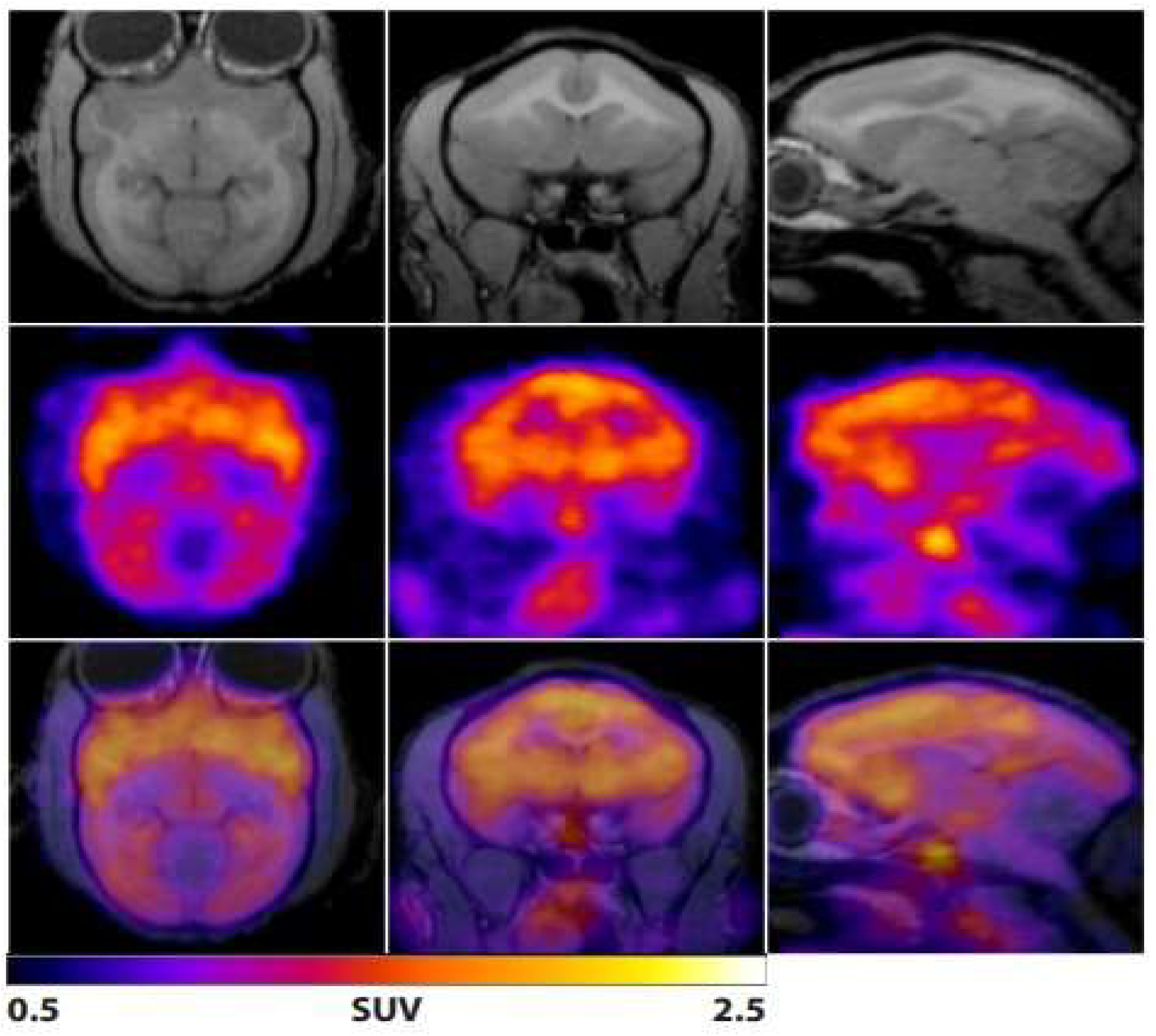
Representative neuroimaging data from a single titi monkey illustrating the image analysis pipeline. Axial (left column), coronal (middle column), and sagittal (right column) views are shown for anatomical T1-weighted MR images (top row), [^11^C]GR103545 standardized uptake value (SUV) maps (middle row), and PET data (color) overlaid on MRI data (grayscale; bottom row). MRI data were bias-corrected and up-sampled in the coronal axis to achieve 0.3 mm isotropic voxels, then aligned to the AC-PC plane. PET data were motion-corrected via rigid registration and manually coregistered to the MRI data. PET data shown here are presented as SUV maps calculated from 50 to 60 minutes post radiotracer injection to highlight radiotracer uptake in the nucleus accumbens.

### Regions of Interest

Regions of interest (ROIs) were selected based on their established roles in pair bonding, separation distress, KOR signaling, and OT modulation. ROIs were classified as either a priori or exploratory based on the strength of prior evidence, which also determined the multiple comparisons correction strategy (see Statistical Analysis). As the effect of separation was not anticipated to be unihemispheric, BP_ND_ values from the left and right hemispheres were combined into a single bilateral estimate per region using a voxel-weighted average:

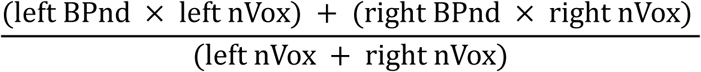

#### A priori regions

A priori regions were selected based on direct evidence for their involvement in pair bonding, social separation, or KOR-OT interactions, and included the hypothalamus, pituitary, NAc, amygdala, lateral septum, hippocampus, and anterior cingulate cortex (ACC).

The hypothalamus and pituitary were selected given their central roles in OT synthesis and release. Bales and Rogers (2022) proposed that during long-term separation, downregulation of KORs in the PVN drives increases in central and peripheral OT, positioning the hypothalamus as a key locus of KOR-OT interaction in the context of partner loss.

The NAc, amygdala, hippocampus, and lateral septum were selected based on their roles in social reward, pair bond maintenance, and the stress response. KORs in the NAc shell have been directly implicated in pair bond maintenance, with KOR antagonist administration in this region attenuating selective aggression toward a stranger in paired male prairie voles (Resendez et al., 2012), and KOR activity in the NAc has been linked to the aversive motivational state of partner loss (Bales & Rogers, 2022). The amygdala has been proposed as a candidate site for oxytocinergic regulation of social behavior and also contains kappa- and mu-opioid receptor-linked pathways, positioning it as a region where KOR signaling and OT-mediated social processes may converge (Putnam & Chang, 2022). In rats, KORs in the basolateral amygdala have been shown to mediate anxiety (Knoll et al., 2011), and stress-induced dysphoria engages the dynorphin/KOR system in the hippocampus, basolateral amygdala, and NAc (Land et al., 2008), suggesting KORs in these regions may similarly be activated during separation-induced anxiety in titi monkeys. The lateral septum was selected based on evidence from titi monkeys specifically showing differences in glucose uptake between paired and unpaired males in this region (Bales et al., 2017), as well as its established role in stress responses and anxiety (Singewald et al., 2011).

The ACC was selected based on its established role in processing the affective component of social pain and separation distress and has garnered interest due to the presence of both oxytocinergic and opioidergic systems (Putnam & Chang, 2022). KOR signaling in this region has been directly implicated in negative affect, with increased KOR activity in the ACC sufficient to produce conditioned place aversion in mice (Lillo Vizin et al., 2024). In humans, lower ACC KOR availability measured using [^11^C]GR103545 PET has been associated with greater depression severity in individuals with major depressive disorder (Smart et al., 2021), supporting the relevance of this region for KOR-mediated affective processing in the present study.

#### Exploratory Regions

Exploratory regions were selected based on their broader involvement in reward, social behavior circuitry, and/or the presence of moderate-to-high KOR expression. These include the orbitofrontal cortex (OFC; medial, lateral, and caudal subdivisions), insula, ventral pallidum, caudate, putamen, claustrum, parahippocampal cortex, and ventral and medial thalamus. These regions broadly participate in networks that integrate stress, emotion, social reward, and motivation. The OFC show high [^11^C]GR103545 binding in non-human primates (Almeida et al., 2025) and has been linked to social reward valuation (Rolls et al., 2020). The insula also shows high KOR binding (Almeida et al., 2025) and is implicated in interoception and social pain (Eisenberger, 2012). The ventral pallidum is a core node in the social reward circuit that receives dynorphinergic input from the NAc and modulates reward-seeking behavior via KOR signaling (Frankel et al., 2008; Q. Sun et al., 2025). The caudate and putamen both have detectable [^11^C]GR103545 binding confirmed in titi monkeys (Almeida et al., 2025). The claustrum had high KOR availability in titi monkeys (Almeida et al., 2025), consistent with studies demonstrating high KOR expression density in rats (Mansour et al., 1987). The parahippocampal cortex was included based on its close anatomical and functional connections with the hippocampus and its role in the encoding and retrieval of socially relevant and contextual memories (Chen et al., 2021; Davies et al., 2022). The ventral and medial thalamus were included based on their roles as key relay hubs for limbic and affective information processing, including the integration of stress-related signals (Zhang et al., 2019), with detectable [^11^C]GR103545 binding confirmed in titi monkeys (Almeida et al., 2025).

### Statistical Analyses

All statistical analyses were conducted in R (version 4.2.2), and the significance threshold was set at α = .05 for all analyses. Linear mixed effects models (LMEs) were used to examine the effects of condition (baseline vs. separation) and sex (female vs. male) on all dependent variables, with subject included as a random intercept to account for the repeated measures within-subject design. Models were fit using the lmer() function from the lme4 and lmerTest packages (Bates et al., 2015; Kuznetsova et al., 2017), with degrees of freedom estimated using the Satterthwaite approximation. Fixed effects were condition, sex, and their interaction. For all models, residuals were inspected for normality using the Shapiro-Wilk test and visually using histograms and quantile-quantile plots. Standardized residuals were inspected for outliers, defined as values exceeding |z| = 3. No outliers were detected in any model. CSF OT was analyzed using the same LME approach, though results should be interpreted in the context of the limited number of valid observations.

Plasma OT values were log-transformed prior to analysis to satisfy the normality assumption of the linear mixed effects model, which was confirmed using the Shapiro-Wilk test on the residuals of the log-transformed model. Estimated marginal means (EMMs) and their standard errors were calculated using the emmeans package and back-transformed to the original scale for reporting. Back-transformed EMMs represent geometric means with back-transformed 95% confidence intervals.

For the plasma OT model, which yielded a significant Condition × Sex interaction, follow-up simple effects tests were conducted to examine the effect of condition within each sex separately, and the effect of sex within each condition separately, using the pairs() function from the emmeans package. No correction for multiple comparisons was applied to these simple effects tests given their role in unpacking a single significant interaction.

For the PET data, a separate LME model was fit for each region of interest with BP_ND_ as the dependent variable. Regions were analyzed using bilateral BP_ND_ values computed as a voxel-weighted average of the left and right hemispheres. No correction for multiple comparisons was applied to the a priori regions given their hypothesis-driven selection. For the exploratory regions, false discovery rate (FDR) correction was applied separately per effect using the Benjamini-Hochberg procedure to control for multiple comparisons across regions within each effect type (condition main effect, sex main effect, and condition × sex interaction).

## RESULTS

### Cortisol

A linear mixed-effects model was used to examine the effects of condition (baseline vs. separation) and sex (female vs. male) on plasma cortisol, with subject included as a random intercept to account for repeated measures. There was a significant main effect of condition, such that cortisol levels were higher during the separation condition compared to the baseline condition (β = 10.84, SE = 2.82, t(14) = 3.84, p = 0.002, d = 1.03). There was no significant main effect of sex (β = 0.40, SE = 2.91, t(28) = 0.14, p = 0.893, d = 0.03), indicating that males and females did not differ in overall cortisol levels. The condition × sex interaction was not significant (β = 6.18, SE = 3.99, t(14) = 1.55, p = 0.143, d = 0.41), suggesting that the increase in cortisol from baseline to separation did not differ significantly between males and females. See Figure 3 for a visualization of cortisol levels across conditions by sex.

**Figure 3.**
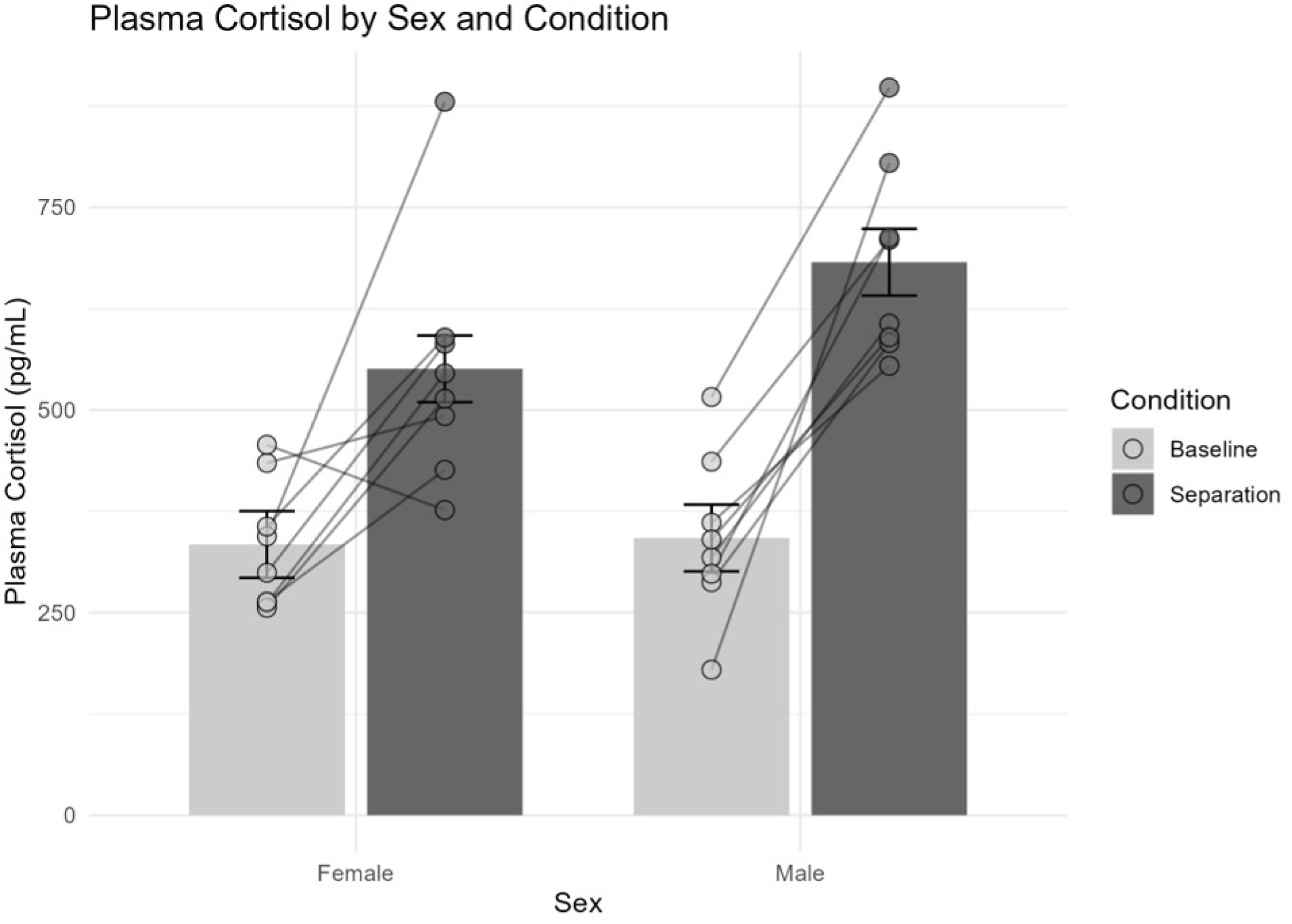
Plasma cortisol levels by condition and sex. Bars represent estimated marginal means derived from the linear mixed effects model, with error bars indicating ± 1 standard error. Individual data points are overlaid. Cortisol was significantly elevated during the separation condition relative to baseline (β = 10.84, SE = 2.82, t(14) = 3.84, p = .002), with no significant main effect of sex and no significant Condition × Sex interaction.

### Oxytocin

A linear mixed effects model was used to examine the effects of condition (baseline vs. separation) and sex (female vs. male) on plasma OT, with subject included as a random intercept to account for repeated measures. Plasma OT values were log-transformed prior to analysis to satisfy the normality assumption of the model, which was confirmed using the Shapiro-Wilk test. There was no significant main effect of condition on log-transformed plasma OT (β = 0.55, SE = 0.35, t(25) = 1.59, p = .124, d = 0.32), indicating that plasma OT did not change significantly from baseline to separation overall. There was a significant main effect of sex (β = 0.70, SE = 0.33, t(25) = 2.10, p = .046, d = 0.42), with males showing higher plasma oxytocin than females overall. Critically, there was a significant Condition × Sex interaction (β = −1.33, SE = 0.48, t(25) = −2.77, p = .010, d = 0.55), indicating that the effect of partner separation on plasma OT differed between males and females. Inspection of the estimated marginal means revealed that females showed a non-significant increase in plasma OT from baseline to separation (baseline geometric mean: 34.2 pg/mL, 95% CI [20.6, 56.8]; separation geometric mean: 59.2 pg/mL, 95% CI [35.6, 98.4]; t(13.6) = −1.57, p = .139, d = 0.43), whereas males showed a significant decrease (baseline geometric mean: 68.9 pg/mL, 95% CI [43.1, 110.3]; separation geometric mean: 31.5 pg/mL, 95% CI [18.9, 52.4]; t(12.6) = 2.33, p = .037, d = 0.66), indicating that the interaction was primarily driven by the male decrease in plasma OT during separation. Follow-up comparisons of males and females within each condition revealed that males had significantly higher plasma OT than females at baseline (t(25) = 2.09, p = .047, d = 0.42), while the sex difference during separation did not reach significance, with females showing a trend toward higher plasma OT than males (t(25) = −1.81, p = .083, d = 0.36). See Figure 4 for a visualization of plasma OT levels across conditions for males and females.

**Figure 4.**
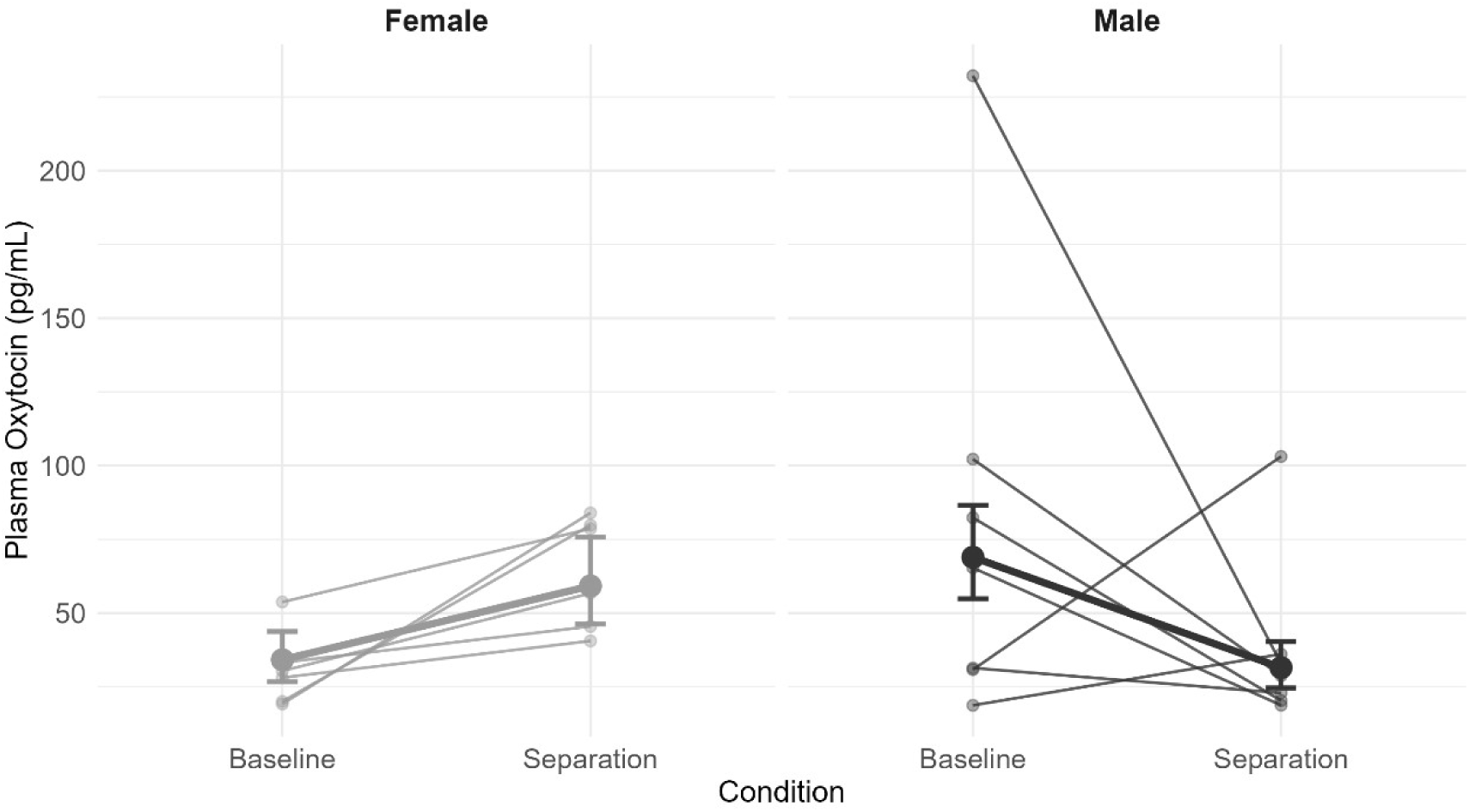
Plasma OT levels by condition and sex. Each panel displays individual subject trajectories (thin lines) alongside estimated marginal means derived from the linear mixed effects model (thick lines), with error bars indicating ± 1 back-transformed standard error. Estimated marginal means represent geometric means on the original scale following back-transformation from the log scale. A significant Condition × Sex interaction was observed (β = −1.33, SE = 0.48, t(25) = −2.77, p = .010), with males showing a significant decrease in plasma oxytocin during separation (t(12.6) = 2.33, p = .037) and females showing a non-significant increase (t(13.6) = 1.57, p = .139). Individual lines represent subjects with complete plasma oxytocin data at both conditions.

**Figure 5.**
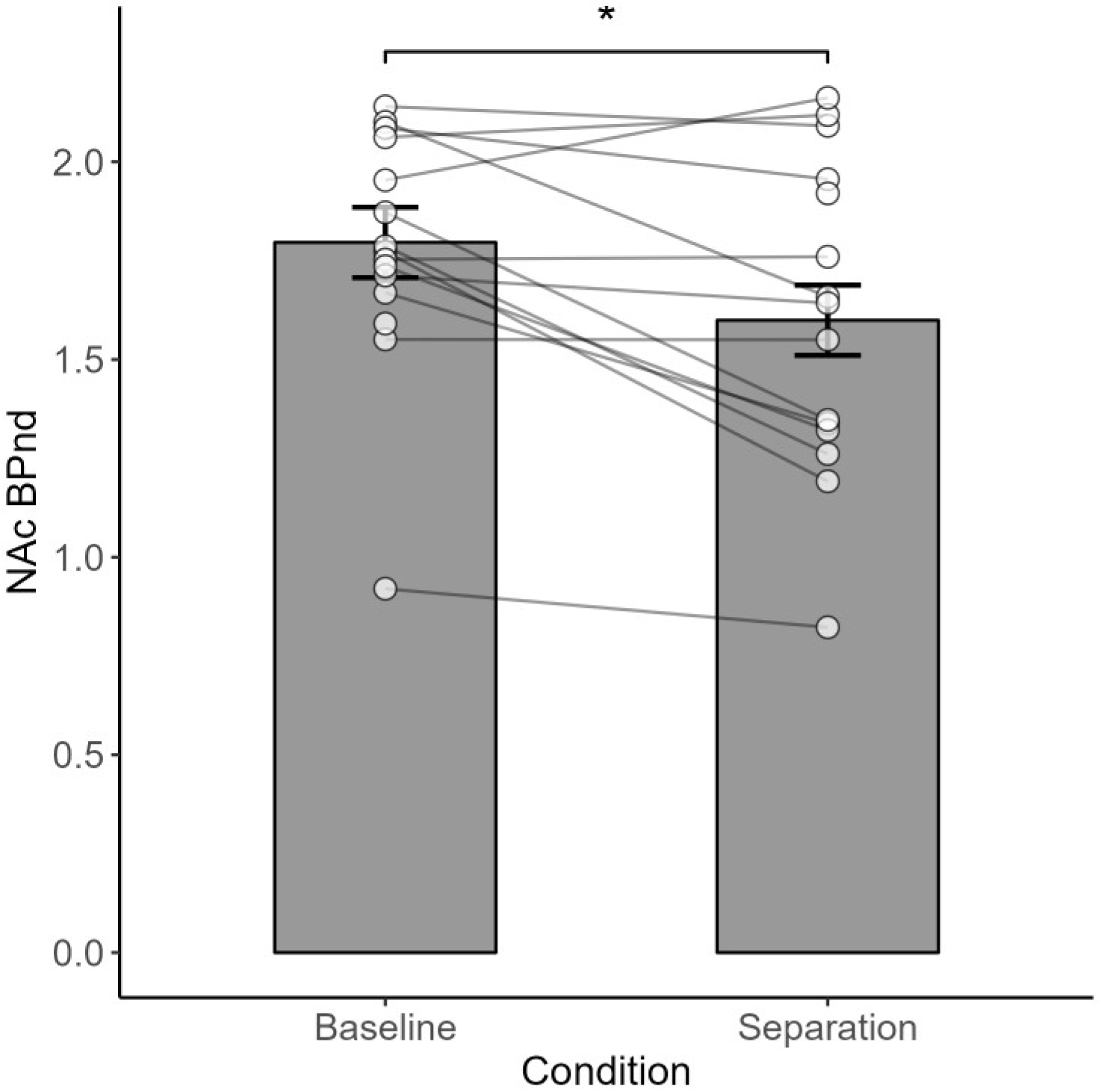
KOR binding potential (BP_ND_) in the nucleus accumbens (NAc) by condition. Bars represent estimated marginal means derived from the linear mixed effects model, collapsed across sex, with error bars indicating ± 1 standard error. Individual data points are overlaid. Partner separation was associated with significantly reduced NAc BP_ND_ relative to baseline (β = −0.28, SE = 0.12, t = −2.99, p = .011), with no significant main effect of sex and no significant Condition × Sex interaction. *p < .05.

A similar linear mixed effects model was used to examine the effects of condition (baseline vs. separation) and sex (female vs. male) on CSF OT, with subject included as a random intercept to account for repeated measures. A substantial proportion of samples were unavailable due to insufficient CSF volume collected during the procedure (n = 11 missing), values falling below the assay detection threshold (n = 2 undetected), or concentrations below the lowest standard of the assay (n = 6 below 16 pg/mL), resulting in only 13 valid observations across 10 subjects.

There was no significant main effect of condition on CSF OT (β = 5.05, SE = 21.28, t(4.21) = 0.24, p = .824, d = 0.12), no significant main effect of sex (β = 20.61, SE = 19.57, t(9.00) = 1.05, p = .320, d = 0.35), and no significant Condition × Sex interaction (β = −22.68, SE = 32.20, t(5.02) = −0.70, p = .513, d = 0.31). These null findings should be interpreted with caution given the limited number of valid observations and the corresponding reduction in statistical power.

### PET imaging

For each region of interest, KOR binding potential (BP_ND_) was analyzed using a linear mixed-effects model with condition (baseline vs. separation) and sex (female vs. male) as fixed effects and subject included as a random intercept. Regions were classified as either *a priori* or exploratory based on their established role in social bonding and separation distress.

#### A Priori Regions

In the *a priori* regions, partner separation was associated with significantly reduced BP_ND_ in the NAc (β = −0.28, SE = 0.12, t(12.75) = −2.99, p = 0.011, d = 0.84), indicating reduced KOR availability for radioligand binding during separation relative to baseline, consistent with either down-regulated KOR or increased endogenous dynorphin occupancy in this region. The ACC showed a trend toward reduced BP_ND_ during separation that did not reach statistical significance (β = −0.27, SE = 0.13, t(12.23) = −2.03, p = .065, d = 0.58). No significant condition effects were observed in the Amygdala, Hippocampus, Lateral Septum, Hypothalamus, or Pituitary. No significant main effects of sex were observed in any a priori region. No significant Condition × Sex interactions were detected in any a priori region. Full results for all a priori regions are presented in Table 1.

**Table 1.** Linear mixed-effects model results for KOR availability (BP_ND_) in a priori regions of interest. Each region was analyzed using a separate model with Condition (baseline vs. separation), Sex (female vs. male), and their interaction as fixed effects, and Subject included as a random intercept to account for repeated measures. Significant p-values (p < .05) are bolded. df = degrees of freedom.

| Region | Effect | Estimate | Std. Error | df | t-value | p-value |
| --- | --- | --- | --- | --- | --- | --- |
| Anterior Cingulate Cortex | Condition | -0.27 | 0.13 | 12.23 | -2.03 | 0.065 |
|  | Sex | 0.12 | 0.20 | 20.81 | 0.62 | 0.539 |
|  | Condition × Sex | 0.06 | 0.19 | 12.23 | 0.29 | 0.774 |
| Hippocampus | Condition | 0.01 | 0.06 | 14.12 | 0.11 | 0.914 |
|  | Sex | 0.12 | 0.07 | 24.14 | 1.77 | 0.089 |
|  | Condition × Sex | -0.04 | 0.08 | 13.74 | -0.56 | 0.586 |
| Lateral Septum | Condition | -0.14 | 0.08 | 13.96 | -1.70 | 0.111 |
|  | Sex | 0.20 | 0.11 | 20.96 | 1.77 | 0.091 |
|  | Condition × Sex | -0.04 | 0.11 | 13.68 | -0.36 | 0.726 |
| Amygdala | Condition | 0.01 | 0.12 | 14.37 | 0.06 | 0.957 |
|  | Sex | 0.21 | 0.12 | 26.36 | 1.72 | 0.096 |
|  | Condition × Sex | 0.00 | 0.16 | 13.94 | 0.03 | 0.977 |
| Nucleus Accumbens | Condition | -0.28 | 0.09 | 12.75 | -2.99 | **0.011** |
|  | Sex | 0.14 | 0.17 | 18.70 | 0.78 | 0.445 |
|  | Condition × Sex | 0.16 | 0.13 | 12.75 | 1.20 | 0.252 |
| Hypothalamus | Condition | -0.01 | 0.25 | 14.38 | -0.05 | 0.961 |
|  | Sex | 0.14 | 0.27 | 26.01 | 0.53 | 0.599 |
|  | Condition × Sex | 0.39 | 0.35 | 13.95 | 1.11 | 0.286 |
| Pituitary | Condition | 0.37 | 0.23 | 14.20 | 1.62 | 0.126 |
|  | Sex | -0.10 | 0.28 | 23.74 | -0.36 | 0.719 |
|  | Condition × Sex | 0.04 | 0.31 | 13.84 | 0.13 | 0.895 |

#### Exploratory Regions

In the exploratory regions, two uncorrected effects were observed; however, neither survived false discovery rate (FDR) correction. A significant Condition × Sex interaction was observed in the Caudal Orbitofrontal Cortex (β = 0.61, SE = 0.25, t = 2.42, p = .032, d = 0.68), and males showed higher BP_ND_ than females in the Putamen (β = 0.31, SE = 0.14, t = 2.12, p = .047, d = 0.49). Given that neither effect survived FDR correction, these findings should be interpreted with caution. Full results for all exploratory regions including FDR-corrected p-values are presented in Supplementary Table 1.

## DISCUSSION

The present study used PET with [^11^C]GR103545 to examine *in vivo* KOR availability during long-term partner separation in a socially monogamous primate species, alongside complementary measures of plasma OT, CSF OT, and plasma cortisol. While recent work has examined KOR availability during acute stress and social buffering in this species (Manca et al., 2026), the present study is the first to directly examine KOR availability during prolonged partner separation using in vivo PET imaging, providing a novel test of the expanded neurobiological model of partner loss proposed by Bales and Rogers (2022).

As a first step, we examined whether the two-week partner separation successfully elicited a physiological stress response, as indexed by plasma cortisol. Cortisol was significantly elevated during separation relative to baseline in both males and females, with no significant sex difference, confirming that the separation paradigm functioned as an effective social stressor and that the HPA axis response was similarly engaged in both sexes. This finding is broadly consistent with evidence that partner separation elevates glucocorticoids in pair-bonded species, including corticosterone elevations following partner loss in prairie voles (Bosch et al., 2009). It is worth noting that Hinde et al. (2016), using a comparable two-week separation paradigm in male titi monkeys, found that plasma cortisol was elevated at 48 hours but no longer significantly elevated at 2 weeks, which differs from the present finding. The reason for this discrepancy is unclear but may reflect individual variation in the duration of HPA activation following partner loss, the larger sample size in the present study, or the inclusion of both sexes in the present study. Regardless, the present finding of elevated cortisol during long-term separation confirms that the two-week separation was physiologically meaningful, and establishes that the neurobiological changes observed in the PET and OT data occurred in the context of a confirmed physiological stress response.

Partner separation was associated with significantly reduced KOR availability in the NAc, providing *in vivo* neuroimaging evidence consistent with endogenous dynorphin release in response to partner loss. Hinde et al. (2016), who examined neural responses to separation in male titi monkeys using [^18^F]-fluorodeoxyglucose ([^18^F]FDG) PET imaging, did not find significant changes in NAc glucose metabolism during either short-term or long-term separation, suggesting that the NAc change observed in the present study may reflect a neurochemical alteration specific to the KOR system that was not detectable with metabolic imaging. This highlights the complementary value of PET imaging with a KOR-targeted radiotracer over generic [^18^F]FDG for characterizing the neurochemical substrates of partner separation.

The decrease in KOR BP_ND_ in the NAc is broadly consistent with the prediction of Bales and Rogers (2022) that KOR activity in the NAc is engaged during partner separation to sustain the aversive motivational state associated with partner loss. However, the precise interpretation of this finding requires careful consideration. Decreased BP_ND_ can result from increased endogenous dynorphin occupancy competing with the radiotracer or receptor downregulation, and these two mechanisms have opposing implications for the model. Bales and Rogers hypothesized that KORs in the NAc would be upregulated during long-term separation, which would manifest as increased rather than decreased BP_ND_. This prediction was explicitly conditional, however, framed as a possibility contingent on CRH levels returning toward baseline over the course of prolonged separation, at which point compensatory KOR upregulation in the NAc would sustain the aversive motivational state in the absence of CRH-driven dynorphin release. The present data suggest that this condition may not yet have been met at the two-week timepoint. Cortisol was significantly elevated during separation relative to baseline, and since cortisol is a downstream product of CRH-driven HPA axis activation, elevated cortisol indirectly suggests that CRH signaling remained elevated at 2 weeks rather than having returned to baseline. If CRH had not yet subsided, the Bales and Rogers prediction of compensatory NAc KOR upregulation would not be expected to have occurred. Under this interpretation, the observed decrease in NAc BP_ND_ most plausibly reflects ongoing dynorphin release driven by continued CRH-mediated stress rather than receptor downregulation, and is therefore consistent with the acute rather than the long-term phase of the Bales and Rogers model.

However, evidence from other stress paradigms suggests that dynorphin/KOR system adaptations during prolonged stress may be more complex than a simple “sustained dynorphin release” interpretation would predict. Studies of chronic social defeat stress in mice have found that while acute stress increases dynorphin mRNA in the NAc, chronic (10-day) social defeat stress produces long-lasting decreases in dynorphin mRNA in the same region (Donahue et al., 2015), suggesting that the dynorphin system may undergo homeostatic downregulation of expression over time rather than remaining hyperactive. Notably, dynorphin adaptations during prolonged stress appear to be region-specific. While chronic social defeat stress decreases dynorphin mRNA in the NAc (Donahue et al., 2015), the same paradigm has been shown to increase dynorphin A expression in the amygdala when extended to 14 days (Zan et al., 2022), suggesting that different brain regions engage the dynorphin/KOR system in distinct ways during prolonged stress. At the receptor level, additional mechanisms have been identified that can produce persistent alterations in KOR function without changes in ligand availability. Repeated stress has been shown to produce functional dysregulation of KOR signaling through a p38α MAPK-dependent mechanism that involves postsynaptic desensitization of KOR-activated GIRK currents rather than changes in receptor expression (Lemos et al., 2012). More recently, a single exposure to acute stress has been shown to induce persistent constitutive activation of KORs in the VTA lasting at least five days, whereby the receptor remains functionally active in the absence of continued dynorphin binding through a shift to a constitutively active conformation (Polter et al., 2017). Taken together, these findings indicate that the dynorphin/KOR system can undergo multiple distinct adaptations following stress exposure, including changes in ligand expression, receptor desensitization, and constitutive receptor activation, each with different time courses and underlying mechanisms. The two-week partner separation examined here is intermediate in duration between the acute and chronic timepoints characterized in the rodent literature, and the specific temporal dynamics of these dynorphin and KOR adaptations during partner separation in a primate model remain uncharacterized. The observed decrease in NAc BP_ND_ could therefore reflect any of several possibilities: ongoing dynorphin release consistent with a prolonged acute stress response, or receptor downregulation and functional desensitization consistent with adaptations observed in other chronic stress paradigms. Additional mechanisms such as constitutive receptor activation may also contribute to persistent KOR system dysregulation during prolonged stress, though whether such adaptations would be detectable with PET imaging of ligand binding remains uncertain. Direct measurement of dynorphin expression and receptor signaling alongside PET imaging would be necessary to distinguish these possibilities in future studies.

It is worth noting, however, that cortisol and CRH do not always track perfectly under chronic stress conditions, as negative feedback mechanisms can dissociate them, and direct measurement of CRH would be necessary to confirm this interpretation. A longer separation duration that allows the HPA axis to normalize may be required to observe the compensatory NAc KOR upregulation predicted by Bales and Rogers, and future studies examining KOR dynamics across multiple separation durations would be valuable in this regard. This interpretation is consistent with evidence from other paradigms of chronic aversive stimulation, where prolonged dynorphin/KOR system engagement contributes to long-term neuroadaptations that produce persistent negative affective states (Heilig et al., 2010; Meinhardt & Sommer, 2015). An alternative interpretation of the NAc finding is that the decreased BP_ND_ reflects KOR downregulation as a desensitization response to prolonged dynorphin activity over the course of the two-week separation. Under this interpretation, the acute phase of separation would produce robust dynorphin release and KOR activation, and the prolonged signaling at these receptors would trigger compensatory downregulation as a homeostatic mechanism. Under this interpretation, the KOR system would still be the substrate generating the aversive motivational state, but a decrease in KOR activity over the course of prolonged separation would result in a gradual reduction of the aversive response, in contrast to the sustained or upregulated KOR activity hypothesized by Bales and Rogers (2022). While this interpretation cannot be ruled out by the present data, it requires assumptions that are not directly supported by independent evidence, including that the KOR system would return toward baseline before the CRH-driven stress response does, and that the persistent cortisol elevation observed here reflects motivational arousal rather than ongoing stress. The simpler interpretation, that decreased NAc BP_ND_ reflects ongoing dynorphin release driven by continued CRH-mediated stress, is more parsimonious and is directly supported by the elevated cortisol observed at the two-week timepoint.

The ACC showed a non-significant trend toward reduced BP_ND_ during separation in the expected direction, consistent with the well-established role of this region in processing the affective component of social pain (Rotge et al., 2015) and with evidence that KOR signaling in the ACC mediates negative affect and aversive states (Lillo Vizin et al., 2024). The absence of a significant ACC effect may reflect limited statistical power given the small sample size, and the directional consistency with predictions suggests this region warrants further investigation in larger samples. Taken together, the NAc and ACC findings suggest that partner separation engages the dynorphin/KOR system in regions central to both the motivational and affective dimensions of social loss, providing the first *in vivo* primate evidence that KOR availability in these regions changes in response to partner separation.

The remaining *a priori* regions, including the amygdala, lateral septum, hypothalamus, and pituitary, did not show significant condition effects, which may reflect limited statistical power, species-specific differences in the time course of KOR adaptation to separation, or genuine null effects in these regions at the two-week timepoint. It is worth noting that Hinde et al. (2016) found reduced glucose metabolism in the central amygdala during long-term separation in male titi monkeys, while the present study found no significant condition effect in the amygdala for KOR availability. Rather than being contradictory, these findings suggest that the reduced amygdala BP_ND_ observed at 2 weeks may be driven by neurochemical systems other than the KOR system, and underscore the importance of modality-specific imaging for characterizing the distinct neurochemical contributions to the separation response.

The null findings in the hypothalamus and pituitary are particularly noteworthy given that Bales and Rogers (2022) predicted KOR downregulation in both regions during long-term separation as a mechanism driving increased OT release. However, several considerations suggest these null results should be interpreted cautiously rather than as straightforward evidence against the model:

First, with respect to the hypothalamus, there is genuine uncertainty about the density of KOR expression in this region in titi monkeys specifically. Autoradiographic mapping of opioid receptor distribution in titi monkeys found dense MOR but not KOR binding in the hypothalamus (Ragen et al., 2015), and a similar pattern has been noted in prairie voles, which also lack high KOR density in the hypothalamus relative to rats and mice (Resendez et al., 2012). This raises the possibility that the hypothalamic KOR system may be less robustly expressed in these species than in the rodent models on which the Bales and Rogers model is primarily based, which would limit the magnitude of any detectable KOR change in this region during separation regardless of whether downregulation occurred. This interpretation is further supported by Almeida et al. (2025), who found evidence of hypothalamic KOR binding in titi monkeys using the Logan reference tissue model but not the Simplified reference tissue model, with the latter providing a better fit to the data overall. Since the present study used the SRTM exclusively, the detection of hypothalamic KOR changes may have been limited by the sensitivity of this approach in a region of potentially sparse KOR expression. Beyond these species-specific and methodological considerations, two additional factors may have contributed to the null hypothalamic finding. First, the PVN is a small, densely packed nucleus, and the hypothalamus ROI used in the present study encompasses the broader hypothalamic region rather than the PVN specifically, such that any KOR changes localized to the PVN could be diluted by signal from surrounding hypothalamic tissue where no change occurred. Second, the neurobiological transition from acute to long-term separation may occur on a longer timescale in primates than in rodents, such that hypothalamic KOR downregulation may not yet be detectable at the two-week timepoint examined here.

The pituitary presents a similar interpretive challenge. Almeida et al. (2025) found significant KOR binding using the SRTM but not the Logan reference tissue model, a pattern opposite to that observed for the hypothalamus. The inconsistency of KOR binding estimates across kinetic modeling approaches in both the hypothalamus and pituitary suggests that KOR signal in these regions may be at or near the lower boundary of reliable detection with current PET methodology in titi monkeys. The pituitary is an exceptionally small structure that is particularly susceptible to partial volume effects, which dilute the measured signal by including signal from surrounding tissue, and its proximity to vascular structures may introduce additional measurement artifacts. Together, these technical challenges may have reduced the sensitivity of the present analysis to detect pituitary KOR changes even if they occurred.

Taken together, the null findings in the hypothalamus and pituitary do not preclude the possibility that KOR downregulation occurs in these regions during long-term partner separation in this species. Rather, they may highlight important species-specific differences in hypothalamic KOR expression relative to the rodent models on which the Bales and Rogers framework is based, and underscore the need for higher-resolution imaging approaches capable of targeting the PVN and pituitary specifically, as well as studies examining KOR dynamics across multiple separation durations in primate models.

In the exploratory regions, two uncorrected effects were observed but neither survived FDR correction. A Condition × Sex interaction was observed in the caudal orbitofrontal cortex, and males showed higher BP_ND_ than females in the putamen. Given that neither effect survived correction for multiple comparisons, these findings should be interpreted with caution and treated as hypothesis-generating rather than as established effects. However, the involvement of these regions in social reward valuation and stress-related motivational processing respectively warrants further investigation in larger samples. The caudal OFC has been implicated in the integration of social and emotional information (Rolls et al., 2020), supporting its relevance to the context of partner separation, while the putamen warrants further investigation given its role in motivated behavior and its KOR expression (Almeida et al., 2025). Both regions may merit closer examination in future studies of partner separation.

Partner separation was associated with a significant Condition × Sex interaction in plasma OT, reflecting a crossover pattern in which males and females showed opposing trajectories across conditions. Follow-up simple effects tests revealed that the male decrease in plasma OT during separation was statistically significant, while the female increase did not reach statistical significance, suggesting that the male response was the primary statistically detectable driver of the interaction. Notably, Hinde et al. (2016) found no significant change in plasma OT during long-term separation. This discrepancy may reflect individual variation in the plasma OT response to separation in this species.

The significant decrease in plasma OT in males during separation is consistent with the theoretical prediction that KOR-mediated suppression of OT release contributes to reduced peripheral OT during partner loss, as the dynorphin/KOR system is known to inhibit OT secretion and KOR agonists have been shown to decrease circulating OT concentrations in rodents (Summy-Long et al., 1990; Van de Heijning et al., 1991; van Wimersma Greidanus et al., 1996). However, the PET findings discussed above complicate a straightforward KOR-mediated suppression interpretation. Although NAc BP_ND_ was significantly reduced during separation, this effect was not sex-specific, and no significant condition effects were observed in the hypothalamus or pituitary, the regions most directly implicated in peripheral OT release. The absence of sex-specific KOR changes in regions directly linked to peripheral OT secretion means that the present data do not provide direct neuroimaging evidence for KOR-mediated suppression of peripheral OT specifically in males. The male OT decrease may therefore reflect a combination of mechanisms. One possibility is that, in this pair-bonded species, male plasma OT during cohabitation may be strongly driven by partner-directed affiliative contact and proximity. Under this interpretation, the male decrease during separation would primarily reflect the loss of partner-stimulated OT release rather than active KOR-mediated suppression, though the two mechanisms are not mutually exclusive. A second possibility is that the sex-dependent OT response could reflect sex differences in CRH signaling rather than sex differences in KOR sensitivity per se. Since CRH stimulates dynorphin release and subsequent KOR-mediated OT suppression (Bruchas et al., 2010; Land et al., 2008), differential CRH responses to separation in males and females could produce the observed OT pattern through the CRH-dynorphin-KOR cascade without requiring sex differences in KOR expression or availability. Direct measurement of CRH in future studies would be necessary to distinguish between these possibilities.

Males showed significantly higher plasma OT than females at baseline, while this difference reversed in direction during separation, though the reversed difference did not reach statistical significance. This crossover pattern is noteworthy given the mixed evidence on sex differences in baseline plasma OT in humans, with some studies finding higher plasma OT in women (Marazziti et al., 2019) and others finding higher plasma OT in men (Weisman et al., 2013). The higher male baseline OT observed in the present study may reflect species-specific differences or the strong coupling between partner-directed affiliative contact and OT release in pair-bonded males. The divergent OT trajectories observed in males and females may reflect sex-specific functional roles of OT, with females showing affiliative and tend-and-befriend responses to stress, and males’ OT levels being more tightly coupled to partner presence and contact. The directional pattern observed in females, though not reaching statistical significance, is broadly consistent with human studies showing that plasma OT may serve as an index of relationship stress in women specifically (Taylor et al., 2006), with women showing more pronounced increases in plasma OT during interpersonal distress than men (Taylor et al., 2010). The significant decrease in male plasma OT during separation in the present study suggests that male OT dynamics in pair-bonded primates are sensitive to partner loss, and the directional difference from the female response may reflect the differential involvement of vasopressin in males (Taylor et al., 2010) or other mechanisms not captured by plasma OT alone. Together, these findings suggest that the kappa opioid system may play a modulatory role in the OT response to partner separation, though the sex-specific nature of this role and its precise neurobiological mechanism remain to be fully elucidated.

Several limitations of the present study should be acknowledged. First, the sample size of 16 subjects (8 per sex) provides limited statistical power, which may have contributed to the absence of significant effects in several a priori regions and the non-significant female OT response. Effect sizes, while generally in the medium to large range, are estimated with considerable uncertainty given the small sample. Second, decreases in BP_ND_ can reflect either increased endogenous dynorphin occupancy competing with the radiotracer or receptor downregulation, and the present study cannot distinguish between these two mechanisms. This ambiguity is inherent to PET displacement imaging with agonist radiotracers and has important implications for interpreting the NAc finding in the context of the Bales and Rogers model, which makes specific predictions about receptor upregulation rather than endogenous ligand occupancy. Third, CSF OT data were severely limited by missing samples, with only 13 valid observations available across 10 subjects, precluding meaningful analysis of central OT dynamics and leaving the central OT response to partner separation in this species uncharacterized in the present study. Fourth, the ROIs used in the present study encompass broad anatomical regions rather than specific nuclei of interest. This is particularly relevant for the hypothalamus, where the Bales and Rogers model specifically predicts KOR changes in the PVN, and for the pituitary, both of which are exceptionally small structures susceptible to partial volume effects. Higher-resolution imaging approaches capable of targeting these subregions specifically would be necessary to adequately test the model’s predictions for these regions.

Fifth, CRH was not directly measured in the present study, limiting the ability to evaluate the proposed mechanistic link between HPA axis activity and KOR-mediated OT suppression. Direct measurement of CRH would be necessary to confirm whether the cortisol elevations observed during separation reflect sustained CRH activity, and to adjudicate between the competing interpretations of the sex-dependent OT response proposed here. Sixth, although the within-subject design controls for any consistent effect of ketamine sedation on KOR binding measurements, the use of ketamine sedation represents an inherent feature of the imaging protocol that could not be avoided. Finally, the present findings were obtained in a single socially monogamous primate species, and their generalizability to other pair-bonded species, including humans, cannot be assumed. As discussed above, KOR distribution in the titi monkey brain differs from that in rodent models in important ways, particularly in the hypothalamus, suggesting that the neurobiological mechanisms underlying partner separation may not be uniform across species. Studies in additional pair-bonded species would be valuable for establishing the generalizability of the present findings and for refining translational models of partner loss.

Taken together, the present findings suggest that NAc KOR engagement during partner separation occurs similarly in both sexes, while the peripheral OT response shows clear sex specificity. The lack of correspondence between the sex-uniform NAc KOR change and the sex-specific plasma OT change suggests that these may reflect partially independent processes, with the KOR system mediating a shared aversive motivational response to separation in both sexes, and the sex-specific OT response reflecting mechanisms operating in parallel to or downstream of NAc KOR engagement. These parallel mechanisms could include sex differences in PVN CRH signaling, sex differences in the contribution of partner-directed affiliative contact to baseline OT release, or sex differences in peripheral OT clearance, none of which can be directly evaluated with the present data. This framing positions the present findings as broadly supporting a role for the KOR system in the neurobiology of partner separation while highlighting the need for future studies that can dissociate the contributions of central KOR engagement from peripheral OT dynamics.

The present study provides the first *in vivo* neuroimaging evidence of KOR system engagement during long-term partner separation in a pair-bonded primate species, directly testing the expanded neurobiological model of partner loss proposed by Bales and Rogers (2022). Reduced KOR availability in the NAc during separation, combined with elevated cortisol and a sex-dependent plasma OT response, is broadly consistent with the Bales and Rogers framework while also revealing patterns that add important nuance and complexity to the model. Together with recent work examining KOR availability during acute stress and social buffering in this species (Manca et al., 2026), the present findings help build a more complete picture of how the KOR system responds to different aspects of pair bond disruption across different timescales.

These findings underscore the importance of including both sexes in studies of pair bond disruption and highlight the need for translational frameworks that account for cross-species variation in KOR distribution and function. Future studies examining KOR dynamics across multiple separation durations and incorporating direct measurement of CRH will be essential for fully characterizing the neurobiological mechanisms underlying the health consequences of social bond disruption.

## Funding

This work was supported by the National Institutes of Health (grant number P51 OD011107, R01 MH125411, S10 OD021715, S10 OD030440, R33 AT012187, U54 NS127758, and the Good Nature Institute).

## Author Contributions (CRediT)

John P. Paulus – investigation, project administration, visualization, formal analysis, data curation, writing – original draft.

Claudia Manca – investigation, visualization, formal analysis, writing – reviewing and editing.

Alita J. D Almeida – software, validation, writing – reviewing and editing.

Anelise Caceres – software, validation, writing – reviewing and editing.

Meghan J. Sosnowski – methodology, resources, supervision, writing – reviewing and editing.

Brad A. Hobson – software, validation, writing – reviewing and editing.

Emilio Ferrer – formal analysis, writing – reviewing and editing.

Abhijit J. Chaudhari – methodology, resources, supervision, funding acquisition, writing – reviewing and editing.

Karen L. Bales – conceptualization, methodology, resources, supervision, visualization, funding acquisition, writing – reviewing and editing.

## Acknowledgments

The authors gratefully acknowledge the members of the Bales lab for their aid with data collection throughout this study. We thank Gitanjali Gnanadesikan for developing and validating the plasma oxytocin extraction protocol for titi monkey samples. We also recognize Jaleh Janatpour, Kevin Theis, and the veterinary and husbandry staff at NBRI for their exceptional animal care and support. Finally, we thank Joshua Waltenburg, Charles Smith, and Sarah Tam of the UC Davis Center for Molecular and Genomic Imaging for their expertise and assistance with PET imaging procedures.

## Notes

### Competing Interest Statement

The authors have declared no competing interest.

## References

Almeida, A. J. D., Hobson, B. A., Caceres, A., Tam, S., Paulus, J. P., Manca, C., Joshi, A. A., Freeman, S. M., Bales, K. L., & Chaudhari, A. J. (2026). A hierarchical brain MRI atlas of the coppery titi monkey (*Plecturocebus cupreus*). NeuroImage, 333, 121921. 10.1016/j.neuroimage.2026.121921

Almeida, A. J. D., Hobson, B. A., Savidge, L. E., Manca, C., Paulus, J. P., Bales, K. L., & Chaudhari, A. J. (2025). Mapping Kappa Opioid Receptor Binding in Titi Monkeys with [11C]GR103545 PET. Molecular Imaging, 24, 15353508251341082. 10.1177/15353508251341082

Avants, B. B., Tustison, N. J., Song, G., Cook, P. A., Klein, A., & Gee, J. C. (2011). A reproducible evaluation of ANTs similarity metric performance in brain image registration. NeuroImage, 54(3), 2033–2044. 10.1016/j.neuroimage.2010.09.025

Bales, K. L., Ardekani, C. S., Baxter, A., Karaskiewicz, C. L., Kuske, J. X., Lau, A. R., Savidge, L. E., Sayler, K. R., & Witczak, L. R. (2021). What is a pair bond? Hormones and Behavior, 136, 105062. 10.1016/j.yhbeh.2021.105062

Bales, K. L., Arias del Razo, R., Conklin, Q. A., Hartman, S., Mayer, H. S., Rogers, F. D., Simmons, T. C., Smith, L. K., Williams, A., Williams, D. R., Witczak, L. R., & Wright, E. C. (2017). Titi Monkeys as a Novel Non-Human Primate Model for the Neurobiology of Pair Bonding. The Yale Journal of Biology and Medicine, 90(3), 373–387.

Bales, K. L., & Rogers, F. D. (2022). Interactions between the κ opioid system, corticotropin-releasing hormone and oxytocin in partner loss. Philosophical Transactions of the Royal Society, 377(1858), 20210061. 10.1098/rstb.2021.0061

Bates, D., Mächler, M., Bolker, B., & Walker, S. (2015). Fitting Linear Mixed-Effects Models Using lme4. Journal of Statistical Software, 67, 1–48. 10.18637/jss.v067.i01

Borrow, A. P., & Cameron, N. M. (2012). The role of oxytocin in mating and pregnancy. Hormones and Behavior, Oxytocin, Vasopressin and Social Behavior, 61(3), 266–276. 10.1016/j.yhbeh.2011.11.001

Bosch, O. J., Dabrowska, J., Modi, M. E., Johnson, Z. V., Keebaugh, A. C., Barrett, C. E., Ahern, T. H., Guo, J., Grinevich, V., Rainnie, D. G., Neumann, I. D., & Young, L. J. (2016). Oxytocin in the nucleus accumbens shell reverses CRFR2-evoked passive stress-coping after partner loss in monogamous male prairie voles. Psychoneuroendocrinology, 64, 66–78. 10.1016/j.psyneuen.2015.11.011

Bosch, O. J., Nair, H. P., Ahern, T. H., Neumann, I. D., & Young, L. J. (2009). The CRF system mediates increased passive stress-coping behavior following the loss of a bonded partner in a monogamous rodent. Neuropsychopharmacology: Official Publication of the American College of Neuropsychopharmacology, 34(6), 1406–1415. 10.1038/npp.2008.154

Bosch, O. J., & Young, L. J. (2018). Oxytocin and Social Relationships: From Attachment to Bond Disruption. Current Topics in Behavioral Neurosciences, 35, 97–117. 10.1007/7854_2017_10

Bower, J. E., Kemeny, M. E., Taylor, S. E., & Fahey, J. L. (1998). Cognitive processing, discovery of meaning, CD4 decline, and AIDS-related mortality among bereaved HIV-seropositive men. Journal of Consulting and Clinical Psychology, 66(6), 979–986. 10.1037//0022-006x.66.6.979

Bruchas, M. R., Land, B. B., & Chavkin, C. (2010). The dynorphin/kappa opioid system as a modulator of stress-induced and pro-addictive behaviors. Brain Research, Neuropeptides in Stress and Addiction, 1314, 44–55. 10.1016/j.brainres.2009.08.062

Callaghan, C. K., Rouine, J., & O’Mara, S. M. (2018). Chapter 3—Potential roles for opioid receptors in motivation and major depressive disorder. In S. O’Mara (Ed.), Progress in Brain Research (Vol. 239, pp. 89–119). Elsevier. 10.1016/bs.pbr.2018.07.009

Carter, C. S., Kenkel, W. M., MacLean, E. L., Wilson, S. R., Perkeybile, A. M., Yee, J. R., Ferris, C. F., Nazarloo, H. P., Porges, S. W., Davis, J. M., Connelly, J. J., & Kingsbury, M. A. (2020). Is Oxytocin “Nature’s Medicine”? Pharmacological Reviews, 72(4), 829–861. 10.1124/pr.120.019398

Chen, H. J., Qi, R., Ke, J., Qiu, J., Xu, Q., Zhang, Z., Zhong, Y., Lu, G. M., & Chen, F. (2021). Altered dynamic parahippocampus functional connectivity in patients with post-traumatic stress disorder. The World Journal of Biological Psychiatry, 22(3), 236–245. 10.1080/15622975.2020.1785006

Dal Monte, O., Noble, P. L., Turchi, J., Cummins, A., & Averbeck, B. B. (2014). CSF and Blood Oxytocin Concentration Changes following Intranasal Delivery in Macaque. PLoS ONE, 9(8), e103677. 10.1371/journal.pone.0103677

Davies, C., Appiah-Kusi, E., Wilson, R., Blest-Hopley, G., Bossong, M. G., Valmaggia, L., Brammer, M., Perez, J., Allen, P., Murray, R. M., McGuire, P., & Bhattacharyya, S. (2022). Altered relationship between cortisol response to social stress and mediotemporal function during fear processing in people at clinical high risk for psychosis: A preliminary report. European Archives of Psychiatry and Clinical Neuroscience, 272(3), 461–475. 10.1007/s00406-021-01318-z

Ditzen, B., Neumann, I. D., Bodenmann, G., von Dawans, B., Turner, R. A., Ehlert, U., & Heinrichs, M. (2007). Effects of different kinds of couple interaction on cortisol and heart rate responses to stress in women. Psychoneuroendocrinology, 32(5), 565–574. 10.1016/j.psyneuen.2007.03.011

Donahue, R. J., Landino, S. M., Golden, S. A., Carroll, F. I., Russo, S. J., & Carlezon, W. A. (2015). Effects of acute and chronic social defeat stress are differentially mediated by the dynorphin/kappa-opioid receptor system. Behavioural Pharmacology, 26(7 0 0), 654–663. 10.1097/FBP.0000000000000155

Donovan, M., Liu, Y., & Wang, Z. (2018). Anxiety-like behavior and neuropeptide receptor expression in male and female prairie voles: The effects of stress and social buffering. Behavioural Brain Research, 342, 70–78. 10.1016/j.bbr.2018.01.015

Dumais, K. M., & Veenema, A. H. (2016). Vasopressin and oxytocin receptor systems in the brain: Sex differences and sex-specific regulation of social behavior. Frontiers in Neuroendocrinology, 40, 1–23. 10.1016/j.yfrne.2015.04.003

Eisenberger, N. I. (2012). The pain of social disconnection: Examining the shared neural underpinnings of physical and social pain. Nature Reviews Neuroscience, 13(6), 421–434. 10.1038/nrn3231

Frankel, P. S., Alburges, M. E., Bush, L., Hanson, G. R., & Kish, S. J. (2008). Striatal and ventral pallidum dynorphin concentrations are markedly increased in human chronic cocaine users. Neuropharmacology, 55(1), 41–46. 10.1016/j.neuropharm.2008.04.019

Gordon, I., Martin, C., Feldman, R., & Leckman, J. F. (2011). Oxytocin and social motivation. Developmental Cognitive Neuroscience, Special Issue on Motivation, 1(4), 471–493. 10.1016/j.dcn.2011.07.007

Grewen, K. M., Anderson, B. J., Girdler, S. S., & Light, K. C. (2003). Warm Partner Contact Is Related to Lower Cardiovascular Reactivity. Behavioral Medicine, 29(3), 123–130. 10.1080/08964280309596065

Heilig, M., Egli, M., Crabbe, J. C., & Becker, H. C. (2010). Acute withdrawal, protracted abstinence and negative affect in alcoholism: Are they linked? Addiction Biology, 15(2), 169–184. 10.1111/j.1369-1600.2009.00194.x

Heinrichs, M., Baumgartner, T., Kirschbaum, C., & Ehlert, U. (2003). Social support and oxytocin interact to suppress cortisol and subjective responses to psychosocial stress. Biological Psychiatry, 54(12), 1389–1398. 10.1016/s0006-3223(03)00465-7

Hinde, K., Muth, C., Maninger, N., Ragen, B. J., Larke, R. H., Jarcho, M. R., Mendoza, S. P., Mason, W. A., Ferrer, E., Cherry, S. R., Fisher-Phelps, M. L., & Bales, K. L. (2016). Challenges to the Pair Bond: Neural and Hormonal Effects of Separation and Reunion in a Monogamous Primate. Frontiers in Behavioral Neuroscience, 10, 221. 10.3389/fnbeh.2016.00221

Holt-Lunstad, J., Smith, T. B., Baker, M., Harris, T., & Stephenson, D. (2015). Loneliness and Social Isolation as Risk Factors for Mortality: A Meta-Analytic Review. Perspectives on Psychological Science, 10(2), 227–237. 10.1177/1745691614568352

Holt-Lunstad, J., Smith, T. B., & Layton, J. B. (2010). Social relationships and mortality risk: A meta-analytic review. PLoS Medicine, 7(7), e1000316. 10.1371/journal.pmed.1000316

Hopf, D., Eckstein, M., Aguilar-Raab, C., Warth, M., & Ditzen, B. (2020). Neuroendocrine mechanisms of grief and bereavement: A systematic review and implications for future interventions. Journal of Neuroendocrinology, 32(8), e12887. 10.1111/jne.12887

Insel, T. R., & Shapiro, L. E. (1992). Oxytocin receptor distribution reflects social organization in monogamous and polygamous voles. Proceedings of the National Academy of Sciences of the United States of America, 89(13), 5981–5985. 10.1073/pnas.89.13.5981

Irwin, M., Daniels, M., Risch, S. C., Bloom, E., & Weiner, H. (1988). Plasma cortisol and natural killer cell activity during bereavement. Biological Psychiatry, 24(2), 173–178. 10.1016/0006-3223(88)90272-7

Kiecolt-Glaser, J. K., Fisher, L. D., Ogrocki, P., Stout, J. C., Speicher, C. E., & Glaser, R. (1987). Marital quality, marital disruption, and immune function. Psychosomatic Medicine, 49(1), 13–34. 10.1097/00006842-198701000-00002

Kiecolt-Glaser, J. K., & Wilson, S. J. (2017). Lovesick: How Couples’ Relationships Influence Health. Annual Review of Clinical Psychology, 13, 421–443. 10.1146/annurev-clinpsy-032816-045111

Kirschbaum, C., Klauer, T., Filipp, S. H., & Hellhammer, D. H. (1995). Sex-specific effects of social support on cortisol and subjective responses to acute psychological stress. Psychosomatic Medicine, 57(1), 23–31. 10.1097/00006842-199501000-00004

Knoll, A. T., Muschamp, J. W., Daws, S. E., Ferguson, D., Dietz, D. M., Meloni, E. G., Carroll, F. I., Nestler, E. J., Konradi, C., & Carlezon, W. A. (2011). Kappa opioid receptor signaling in the basolateral amygdala regulates conditioned fear and anxiety in rats. Biological Psychiatry, 70(5), 425–433. 10.1016/j.biopsych.2011.03.017

Kuznetsova, A., Brockhoff, P. B., & Christensen, R. H. B. (2017). lmerTest Package: Tests in Linear Mixed Effects Models. Journal of Statistical Software, 82, 1–26. 10.18637/jss.v082.i13

Land, B. B., Bruchas, M. R., Lemos, J. C., Xu, M., Melief, E. J., & Chavkin, C. (2008). The Dysphoric Component of Stress Is Encoded by Activation of the Dynorphin κ-Opioid System. The Journal of Neuroscience, 28(2), 407–414. 10.1523/JNEUROSCI.4458-07.2008

Lemos, J. C., Roth, C. A., Messinger, D. I., Gill, H. K., Phillips, P. E. M., & Chavkin, C. (2012). Repeated Stress Dysregulates κ-Opioid Receptor Signaling in the Dorsal Raphe through a p38α MAPK-Dependent Mechanism. The Journal of Neuroscience, 32(36), 12325– 12336. 10.1523/JNEUROSCI.2053-12.2012

Lillo Vizin, R. C., Ito, H., Kopruszinski, C. M., Ikegami, M., Ikegami, D., Yue, X., Navratilova, E., Moutal, A., Cowen, S. L., & Porreca, F. (2024). Cortical kappa opioid receptors integrate negative affect and sleep disturbance. Translational Psychiatry, 14(1), 417. 10.1038/s41398-024-03123-3

Liu, Y., & Wang, Z. X. (2003). Nucleus accumbens oxytocin and dopamine interact to regulate pair bond formation in female prairie voles. Neuroscience, 121(3), 537–544. 10.1016/s0306-4522(03)00555-4

Loth, M. K., & Donaldson, Z. R. (2020). Oxytocin, Dopamine, and Opioid Interactions Underlying Pair Bonding: Highlighting a Potential Role for Microglia. Endocrinology, 162(2), bqaa223. 10.1210/endocr/bqaa223

Love, T. M. (2018). The impact of oxytocin on stress: The role of sex. Current Opinion in Behavioral Sciences, 23, 136–142. 10.1016/j.cobeha.2018.06.018

Manca, C., Paulus, J. P., Almeida, A. J. D., Caceres, A., Sosnowski, M. J., Hobson, B. A., Ferrer, E., Chaudhari, A. J., & Bales, K. L. (2026). Brain kappa opioid receptor availability across stress and social buffering conditions: A positron emission tomography study in coppery titi monkeys. Neuroscience, 611, 155–169. 10.1016/j.neuroscience.2026.06.028

Mansour, A., Khachaturian, H., Lewis, M. E., Akil, H., & Watson, S. J. (1987). Autoradiographic differentiation of mu, delta, and kappa opioid receptors in the rat forebrain and midbrain. The Journal of Neuroscience: The Official Journal of the Society for Neuroscience, 7(8), 2445–2464.

Marazziti, D., Baroni, S., Mucci, F., Piccinni, A., Moroni, I., Giannaccini, G., Carmassi, C., Massimetti, E., & Dell’Osso, L. (2019). Sex-Related Differences in Plasma Oxytocin Levels in Humans. Clinical Practice and Epidemiology in Mental Health : CP & EMH, 15, 58–63. 10.2174/1745017901915010058

Mason, W. A., & Mendoza, S. P. (1998). Generic aspects of primate attachments: Parents, offspring and mates. Psychoneuroendocrinology, 23(8), 765–778. 10.1016/s0306-4530(98)00054-7

McLaughlin, J. P., Li, S., Valdez, J., Chavkin, T. A., & Chavkin, C. (2006). Social defeat stress-induced behavioral responses are mediated by the endogenous kappa opioid system. Neuropsychopharmacology: Official Publication of the American College of Neuropsychopharmacology, 31(6), 1241–1248. 10.1038/sj.npp.1300872

McNeal, N., Scotti, M.-A. L., Wardwell, J., Chandler, D. L., Bates, S. L., Larocca, M., Trahanas, D. M., & Grippo, A. J. (2014). Disruption of social bonds induces behavioral and physiological dysregulation in male and female prairie voles. Autonomic Neuroscience: Basic & Clinical, 180, 9–16. 10.1016/j.autneu.2013.10.001

Meinhardt, M. W., & Sommer, W. H. (2015). Postdependent state in rats as a model for medication development in alcoholism. Addiction Biology, 20(1), 1–21. 10.1111/adb.12187

Mendoza, S. P., & Mason, W. A. (1986). Contrasting responses to intruders and to involuntary separation by monogamous and polygynous New World monkeys. Physiology & Behavior, 38(6), 795–801. 10.1016/0031-9384(86)90045-4

Nabulsi, N. B., Zheng, M.-Q., Ropchan, J., Labaree, D., Ding, Y.-S., Blumberg, L., & Huang, Y. (2011). [11C]GR103545: Novel one-pot radiosynthesis with high specific activity. Nuclear Medicine and Biology, 38(2), 215–221. 10.1016/j.nucmedbio.2010.08.014

Naganawa, M., Jacobsen, L. K., Zheng, M.-Q., Lin, S.-F., Banerjee, A., Byon, W., Weinzimmer, D., Tomasi, G., Nabulsi, N., Grimwood, S., Badura, L. L., Carson, R. E., McCarthy, T. J., & Huang, Y. (2014). Evaluation of the Agonist PET Radioligand [11C]GR103545 to Image Kappa Opioid Receptor in Humans: Kinetic Model Selection, Test-Retest Reproducibility and Receptor Occupancy by the Antagonist PF-04455242. NeuroImage, 99, 69–79. 10.1016/j.neuroimage.2014.05.033

Panksepp, J., Herman, B. H., Vilberg, T., Bishop, P., & DeEskinazi, F. G. (1980). Endogenous opioids and social behavior. Neuroscience and Biobehavioral Reviews, 4(4), 473–487. 10.1016/0149-7634(80)90036-6

Parker, K. J., Hoffman, C. L., Hyde, S. A., Cummings, C. S., & Maestripieri, D. (2010). Effects of age on cerebrospinal fluid oxytocin levels in free-ranging adult female and infant rhesus macaques. Behavioral Neuroscience, 124(3), 10.1037/a0019576. https://doi.org/10.1037/a0019576

Pohl, T. T., Young, L. J., & Bosch, O. J. (2019). Lost connections: Oxytocin and the neural, physiological, and behavioral consequences of disrupted relationships. International Journal of Psychophysiology: Official Journal of the International Organization of Psychophysiology, 136, 54–63. 10.1016/j.ijpsycho.2017.12.011

Polter, A. M., Barcomb, K., Chen, R. W., Dingess, P. M., Graziane, N. M., Brown, T. E., & Kauer, J. A. (2017). Constitutive activation of kappa opioid receptors at ventral tegmental area inhibitory synapses following acute stress. eLife, 6, e23785. 10.7554/eLife.23785

Putnam, P. T., & Chang, S. W. C. (2022). Interplay between the oxytocin and opioid systems in regulating social behaviour. Philosophical Transactions of the Royal Society of London. Series B, Biological Sciences, 377(1858), 20210050. 10.1098/rstb.2021.0050

Ragen, B. J., Freeman, S. M., Laredo, S. A., Mendoza, S. P., & Bales, K. L. (2015). μ and κ opioid receptor distribution in the monogamous titi monkey (Callicebus cupreus): Implications for social behavior and endocrine functioning. Neuroscience, 290, 421–434. 10.1016/j.neuroscience.2015.01.023

Ragen, B. J., Maninger, N., Mendoza, S. P., Jarcho, M. R., & Bales, K. L. (2013). Presence of a pair-mate regulates the behavioral and physiological effects of opioid manipulation in the monogamous titi monkey (Callicebus cupreus). Psychoneuroendocrinology, 38(11), 10.1016/j.psyneuen.2013.05.009. https://doi.org/10.1016/j.psyneuen.2013.05.009

Remage-Healey, L., Adkins-Regan, E., & Romero, L. M. (2003). Behavioral and adrenocortical responses to mate separation and reunion in the zebra finch. Hormones and Behavior, 43(1), 108–114. 10.1016/S0018-506X(02)00012-0

Resendez, S. L., Dome, M., Gormley, G., Franco, D., Nevárez, N., Hamid, A. A., & Aragona, B. J. (2013). μ-Opioid receptors within subregions of the striatum mediate pair bond formation through parallel yet distinct reward mechanisms. The Journal of Neuroscience: The Official Journal of the Society for Neuroscience, 33(21), 9140–9149. 10.1523/JNEUROSCI.4123-12.2013

Resendez, S. L., Keyes, P. C., Day, J. J., Hambro, C., Austin, C. J., Maina, F. K., Eidson, L. N., Porter-Stransky, K. A., Nevárez, N., McLean, J. W., Kuhnmuench, M. A., Murphy, A. Z., Mathews, T. A., & Aragona, B. J. (2016). Dopamine and opioid systems interact within the nucleus accumbens to maintain monogamous pair bonds. eLife, 5, e15325. 10.7554/eLife.15325

Resendez, S. L., Kuhnmuench, M., Krzywosinski, T., & Aragona, B. J. (2012). κ-Opioid receptors within the nucleus accumbens shell mediate pair bond maintenance. The Journal of Neuroscience: The Official Journal of the Society for Neuroscience, 32(20), 6771–6784. 10.1523/JNEUROSCI.5779-11.2012

Robles, T. F., Slatcher, R. B., Trombello, J. M., & McGinn, M. M. (2014). Marital quality and health: A meta-analytic review. Psychological Bulletin, 140(1), 140–187. 10.1037/a0031859

Rolls, E. T., Cheng, W., & Feng, J. (2020). The orbitofrontal cortex: Reward, emotion and depression. Brain Communications, 2(2), fcaa196. 10.1093/braincomms/fcaa196

Rotge, J.-Y., Lemogne, C., Hinfray, S., Huguet, P., Grynszpan, O., Tartour, E., George, N., & Fossati, P. (2015). A meta-analysis of the anterior cingulate contribution to social pain. Social Cognitive and Affective Neuroscience, 10(1), 19–27. 10.1093/scan/nsu110

Seiler, A., von Känel, R., & Slavich, G. M. (2020). The Psychobiology of Bereavement and Health: A Conceptual Review From the Perspective of Social Signal Transduction Theory of Depression. Frontiers in Psychiatry, 11, 565239. 10.3389/fpsyt.2020.565239

Singewald, G. M., Rjabokon, A., Singewald, N., & Ebner, K. (2011). The Modulatory Role of the Lateral Septum on Neuroendocrine and Behavioral Stress Responses. Neuropsychopharmacology, 36(4), 793–804. 10.1038/npp.2010.213

Smart, K., Yttredahl, A., Oquendo, M. A., Mann, J. J., Hillmer, A. T., Carson, R. E., & Miller, J. M. (2021). Data-driven analysis of kappa opioid receptor binding in major depressive disorder measured by positron emission tomography. Translational Psychiatry, 11(1), 602. 10.1038/s41398-021-01729-5

Smith, A. S., & Wang, Z. (2014). Hypothalamic Oxytocin Mediates Social Buffering of the Stress Response. Biological Psychiatry, Neurobiological Moderators of Stress Response, 76(4), 281–288. 10.1016/j.biopsych.2013.09.017

Smith, T. E., McGreer-Whitworth, B., & French, J. A. (1998). Close proximity of the heterosexual partner reduces the physiological and behavioral consequences of novel-cage housing in black tufted-ear marmosets (Callithrix kuhli). Hormones and Behavior, 34(3), 211–222. 10.1006/hbeh.1998.1469

Summy-Long, J. Y., Rosella-Dampman, L. M., McLemore, G. L., & Koehler, E. (1990). Kappa opiate receptors inhibit release of oxytocin from the magnocellular system during dehydration. Neuroendocrinology, 51(4), 376–384. 10.1159/000125364

Sun, P., Smith, A., Lei, K., Liu, Y., & Wang, Z. (2014). Breaking bonds in male prairie vole: Long-term effects on emotional and social behavior, physiology, and neurochemistry. Behavioural Brain Research, 265, 22–31. 10.1016/j.bbr.2014.02.016

Sun, Q., Liu, M., Guan, W., Xiao, X., Dong, C., Bruchas, M. R., Zweifel, L. S., Li, Y., Tian, L., & Li, B. (2025). Dynorphin modulates reward-seeking actions through a pallido-amygdala cholinergic circuit. Neuron, 113(11), 1823–1840.e8. 10.1016/j.neuron.2025.03.018

Szeto, A., McCabe, P. M., Nation, D. A., Tabak, B. A., Rossetti, M. A., McCullough, M. E., Schneiderman, N., & Mendez, A. J. (2011). Evaluation of enzyme immunoassay and radioimmunoassay methods for the measurement of plasma oxytocin. Psychosomatic Medicine, 73(5), 393–400. 10.1097/PSY.0b013e31821df0c2

Tabak, B. A., Leng, G., Szeto, A., Parker, K. J., Verbalis, J. G., Ziegler, T. E., Lee, M. R., Neumann, I. D., & Mendez, A. J. (2023). Advances in human oxytocin measurement: Challenges and proposed solutions. Molecular Psychiatry, 28(1), 127–140. 10.1038/s41380-022-01719-z

Taylor, S. E., Gonzaga, G. C., Klein, L. C., Hu, P., Greendale, G. A., & Seeman, T. E. (2006). Relation of oxytocin to psychological stress responses and hypothalamic-pituitary-adrenocortical axis activity in older women. Psychosomatic Medicine, 68(2), 238–245. 10.1097/01.psy.0000203242.95990.74

Taylor, S. E., Klein, L. C., Lewis, B. P., Gruenewald, T. L., Gurung, R. A., & Updegraff, J. A. (2000). Biobehavioral responses to stress in females: Tend-and-befriend, not fight-or-flight. Psychological Review, 107(3), 411–429. 10.1037/0033-295x.107.3.411

Taylor, S. E., Saphire-Bernstein, S., & Seeman, T. E. (2010). Are plasma oxytocin in women and plasma vasopressin in men biomarkers of distressed pair-bond relationships? Psychological Science, 21(1), 3–7. 10.1177/0956797609356507

Van de Heijning, B. J. M., -Van den Herik, I. K., & Van Wimersma Greidanus, T. B. (1991). The opioid receptor subtypes μ and κ, but not δ, are involved in the control of the vasopressin and oxytocin release in the rat. European Journal of Pharmacology, 209(3), 199–206. 10.1016/0014-2999(91)90170-U

van Wimersma Greidanus, Tj. B., Janssen, S., Frankhuijzen-Sierevogel, J. C., Maigret, C., & van de Heijning, B. J. M. (1996). Effect of central administration of the κ-opiate receptor agonist U 69.593 on neurohypophyseal hormone levels in blood and cerebrospinal fluid. Neuropeptides, 30(5), 452–455. 10.1016/S0143-4179(96)90009-8

Vijay, A., Wang, S., Worhunsky, P., Zheng, M.-Q., Nabulsi, N., Ropchan, J., Krishnan-Sarin, S., Huang, Y., & Morris, E. D. (2016). PET imaging reveals sex differences in kappa opioid receptor availability in humans, in vivo. American Journal of Nuclear Medicine and Molecular Imaging, 6(4), 205–214.

Weisman, O., Zagoory-Sharon, O., Schneiderman, I., Gordon, I., & Feldman, R. (2013). Plasma oxytocin distributions in a large cohort of women and men and their gender-specific associations with anxiety. Psychoneuroendocrinology, 38(5), 694–701. 10.1016/j.psyneuen.2012.08.011

Williams, A. V., Laman-Maharg, A., Armstrong, C. V., Ramos-Maciel, S., Minie, V. A., & Trainor, B. C. (2018). Acute inhibition of kappa opioid receptors before stress blocks depression-like behaviors in California mice. Progress in Neuro-Psychopharmacology & Biological Psychiatry, 86, 166–174. 10.1016/j.pnpbp.2018.06.001

Witczak, L. R., Arias del Razo, R., Baxter, A., Conley, A. J., Cotterman, R., Dufek, M., Goetze, L. R., Lau, A. R., Mendoza, S. P., Savidge, L. E., & Bales, K. L. (2021). Relationships between cortisol and urinary androgens in female titi monkeys (Plecturocebus cupreus). General and Comparative Endocrinology, 314, 113927. 10.1016/j.ygcen.2021.113927

Wright, E. C., Parks, T. V., Alexander, J. O., Supra, R., & Trainor, B. C. (2018). Activation of Kappa Opioid Receptors in the Dorsal Raphe Have Sex Dependent Effects on Social Behavior in California Mice. Behavioural Brain Research, 351, 83–92. 10.1016/j.bbr.2018.05.011

Zan, G., Sun, X., Wang, Y., Liu, R., Wang, C., Du, W., Guo, L., Chai, J., Li, Q., Liu, Z., & Liu, J. (2022). Amygdala dynorphin/κ opioid receptor system modulates depressive-like behavior in mice following chronic social defeat stress. Acta Pharmacologica Sinica, 43(3), 577–587. 10.1038/s41401-021-00677-6

Zhang, X., Li, X., Steffens, D. C., Guo, H., & Wang, L. (2019). Dynamic changes in thalamic connectivity following stress and its association with future depression severity. Brain and Behavior, 9(12), e01445. 10.1002/brb3.1445

