## Supplementary Material for "Kappa Opioid–Oxytocin Interactions During Long-Term Partner Separation: Insights from PET Imaging in Titi Monkeys (Plecturocebus cupreus)"

| **Region** | **Effect** | **Estimate** | **Std. Error** | **df** | **t-value** | **p-value** | **FDR p-value** |
| --- | --- | --- | --- | --- | --- | --- | --- |
| Medial Orbitofrontal Cortex | Condition | -0.03 | 0.18 | 13.76 | -0.16 | 0.872 | 0.872 |
|  | Sex | -0.28 | 0.20 | 25.66 | -1.45 | 0.159 | 0.457 |
|  | Condition × Sex | 0.08 | 0.26 | 13.76 | 0.32 | 0.756 | 0.793 |
| Lateral Orbitofrontal Cortex | Condition | -0.18 | 0.14 | 12.64 | -1.26 | 0.230 | 0.405 |
|  | Sex | -0.17 | 0.19 | 22.22 | -0.90 | 0.380 | 0.576 |
|  | Condition × Sex | 0.20 | 0.20 | 12.64 | 1.02 | 0.325 | 0.793 |
| Caudal Orbitofrontal Cortex | Condition | -0.28 | 0.18 | 12.67 | -1.56 | 0.143 | 0.398 |
|  | Sex | -0.35 | 0.24 | 22.73 | -1.47 | 0.154 | 0.457 |
|  | Condition × Sex | 0.61 | 0.25 | 12.67 | 2.42 | 0.032 | 0.352 |
| Parahippocampal Cortex | Condition | 0.04 | 0.12 | 14.33 | 0.32 | 0.754 | 0.829 |
|  | Sex | 0.15 | 0.14 | 25.56 | 1.09 | 0.284 | 0.576 |
|  | Condition × Sex | -0.12 | 0.17 | 13.91 | -0.69 | 0.503 | 0.793 |
| Insula | Condition | -0.21 | 0.10 | 13.79 | -2.11 | 0.054 | 0.398 |
|  | Sex | 0.05 | 0.16 | 19.92 | 0.35 | 0.733 | 0.806 |
|  | Condition × Sex | 0.12 | 0.14 | 13.54 | 0.88 | 0.396 | 0.793 |
| Claustrum | Condition | -0.20 | 0.14 | 12.83 | -1.45 | 0.170 | 0.398 |
|  | Sex | 0.11 | 0.21 | 21.06 | 0.52 | 0.610 | 0.746 |
|  | Condition × Sex | 0.10 | 0.19 | 12.83 | 0.50 | 0.628 | 0.793 |
| Ventral Pallidum | Condition | -0.21 | 0.15 | 11.24 | -1.43 | 0.181 | 0.398 |
|  | Sex | 0.03 | 0.22 | 19.58 | 0.11 | 0.911 | 0.911 |
|  | Condition × Sex | 0.36 | 0.22 | 11.28 | 1.68 | 0.121 | 0.665 |
| Caudate | Condition | -0.08 | 0.07 | 13.60 | -1.18 | 0.258 | 0.405 |
|  | Sex | 0.16 | 0.11 | 19.44 | 1.44 | 0.166 | 0.457 |
|  | Condition × Sex | -0.03 | 0.09 | 13.36 | -0.27 | 0.793 | 0.793 |
| Putamen | Condition | -0.14 | 0.08 | 13.70 | -1.66 | 0.119 | 0.398 |
|  | Sex | 0.31 | 0.14 | 18.83 | 2.12 | 0.047 | 0.457 |
|  | Condition × Sex | -0.05 | 0.12 | 13.48 | -0.43 | 0.671 | 0.793 |
| Ventral Thalamus | Condition | -0.02 | 0.04 | 13.56 | -0.52 | 0.615 | 0.802 |
|  | Sex | 0.04 | 0.05 | 21.52 | 0.85 | 0.405 | 0.576 |
|  | Condition × Sex | -0.02 | 0.05 | 13.25 | -0.44 | 0.664 | 0.793 |
| Medial Thalamus | Condition | 0.02 | 0.05 | 27.00 | 0.45 | 0.656 | 0.802 |
|  | Sex | 0.04 | 0.05 | 27.00 | 0.82 | 0.419 | 0.576 |
|  | Condition × Sex | -0.07 | 0.07 | 27.00 | -1.02 | 0.317 | 0.793 |

***Supplementary Table 1. Linear mixed-effects model results for KOR availability (BP_ND_) in exploratory regions of interest.*** *Each region was analyzed using a separate model with Condition (baseline vs. separation), Sex (female vs. male), and their interaction as fixed effects, and Subject included as a random intercept to account for repeated measures. df = degrees of freedom.*

**Titi Monkey Plasma R/A Protocol – 8/2/22**

**Reagents:** Prepare first. Keep reagents in foil (light sensitive) at +4 °C until just before needed.

1) 50 mM Tris-HCl (121.14 g/mol), pH 8.0 (stable at +4 °C for ~ 18 months).
   a. On weigh paper, weigh out 1.393 g Trizma base ([Sigma Aldrich T1503](https://www.sigmaaldrich.com/US/en/product/sigma/t1503)).
   b. Dilute Trizma from (a) into 230 μl DI water.
   c. Measure pH (expect ~10.5):
   d. Add increments of HCl to reach pH 8.0 (±0.03):
      i. You can use any leftover stop solution from Arbor Assays, if you have it. It’s 1 M HCl.
      ii. Expect to add 6-6.2 ml.
      iii. Start by adding 3 ml, then use smaller and smaller steps.
   e. Notes:
   f. Final pH:
2) Dithiothreitol (DTT), 154.253 g/mol (make fresh daily).
   a. We use [Sigma Aldrich D0632](https://www.sigmaaldrich.com/US/en/product/sial/d0632).
   b. In 1.5 ml tube, 125 mM: weigh 0.014 g DTT and add 750 μl DI water.
   c. In 1.5 ml tube, 1.25 mM: dilute 10 μl of 125 mM DTT into 990 μl DI water.
3) N-ethylmaleimide (NEM), 125.13 g/mol (make fresh daily)
   a. We use [Sigma Aldrich E3876](https://www.sigmaaldrich.com/US/en/product/sial/e3876?context=product).
   b. In 1.5 ml tube 125 mM: weigh 0.012 g NEM and add 750 μl DI water.
   c. In 2 ml tube, 1.25 mM: dilute 15 μl of 125 mM NEM into 1485 μl DI water.
4) 80% Acetonitrile (ACN) in H_2_O (v/v). Need >204 ml (stable for ~ 1 year).
   a. We use [HPLC grade](https://www.fishersci.com/shop/products/acetonitrile-hplc-fisher-chemical-10/A9984), which might be overkill (it’s more expensive), but I don’t actually know and haven’t tested it.
   b. In 250 μl bottle, dilute 200 μl of ACN into 50 μl of DI water.
   c. Put at +4 °C until needed.

**Notes:**

- You will only be using the 1.25 mM DTT and NEM for the procedure, however, you need to start at 125 mM to be able to weigh the powders accurately.
- You can adjust these amounts as necessary for different numbers of samples.
- Polypropylene is the best tube material for working with oxytocin (doesn’t adsorb OT as much as glass or polystyrene, supposedly).
- Handle all chemicals in a hood if possible (DTT, in particular smells quite bad, and both DTT and NEM are pretty highly toxic).

**Preparation:**

1) Prepare reagents.
2) Set incubator to 37°C.
3) Thaw samples.
4) Label tubes for sample prep.
5) Sample prep: in 5-ml polypropylene tubes, dilute samples into double volume of Tris-HCl (50 mM, pH 8).
   a. If you have enough volume, I suggest 300 μl of plasma into 600 μl of Tris, based on the parallelism and spike recovery results we have so far.
   b. You can also experiment with smaller volumes, just scaling the Tris (and leaving everything else the same), without much concern. The parallelism looked good down to 0.47x (~125 μl of plasma), although it read towards the bottom of the assay curve, so you may run into a large number of samples measuring below the limit of detection, the lower you go.
   c. It is possible that slightly larger volumes would also work, but that would need to be validated (especially spike recovery at an increased concentration factor). Based on the parallelism, we expect problems at 2x or higher. I’m happy to chat about this, if helpful.
   d. If you can assay all your samples (or at least all of your contrasts, e.g. before and after) at the same dilution, that is ideal.

**Reduction/Alkylation:** Handle DTT, NEM, and ACN in a hood if possible.

6) Reduction (break disulfide bonds):
   a. Add 10 μl of 1.25 mM DTT to each sample. (Can use a repeater pipette.)
   b. Vortex for 30 s.
   c. Incubate in the dark at 37 °C for 45 minutes.
7) Cool samples and incubator to room temperature (~10 minutes).
8) Alkylation (alkyl group binds to sulfur):
   a. Add 30 μl of 1.25 mM NEM to each sample. (Can use a repeater pipette.)
   b. Vortex for 30 s.
   c. Incubate at 22 °C in the dark for 20 minutes.
9) Protein precipitation (get rid of potentially interfering proteins):
   a. Add 1.5 ml ice cold 80% ACN to each tube. (Can use a repeater pipette.)
      i. You should see the sample fizz and turn cloudy—that’s the protein!
   b. Vortex for 30 s.
      i. Make sure all samples are well vortexed, otherwise the whole sample will not precipitate properly.
      ii. Invert by hand a couple of times if necessary.
   c. Centrifuge samples at 3,000 rpm for 15 minutes in the large (Sorvall ST 16) centrifuge. (RCF_max_ = ~2,000 x g. Higher is fine, if the tubes are rated for it!)
   d. While samples are centrifuging, label 5-ml polypropylene tubes for sample supernatants.
   e. Transfer supernatant from original tube to matching new tube.
   f. Discard original tubes (with protein pellets at the bottom).
10) Put samples at -80 °C overnight for lyophilization the next day.
    a. Alternatively, if lyophilization is not available, evaporate immediately in water bath at 37 °C, with compressed or pumped air.
